# The Copper Responsive Transcription Factor SPL7 Represses Key Abscisic Acid Biosynthetic Genes to Balance Growth and Drought Tolerance

**DOI:** 10.1101/2021.06.03.446925

**Authors:** Yanzhi Yang, Jianmei Du, Yihan Tao, Chen Hao, Zheng Kuang, Zhan Li, Lei Li

**Affiliations:** State Key Laboratory of Protein and Plant Gene Research, School of Life Sciences and School of Advanced Agricultural Sciences, Peking University, Beijing 100871, China; Peking-Tsinghua Center for Life Sciences, Academy for Advanced Interdisciplinary Studies, Peking University, Beijing 100871, China

## Abstract

Plants adapt to adverse environments by turning on defense against abiotic stresses, which is mainly orchestrated by the phytohormone abscisic acid (ABA). But how ABA homeostasis is modulated to balance growth and stress responses is still largely unknown. Here we report that prior treatment of Arabidopsis seedling with high copper retardates growth but enhances draught tolerance at later stages by modulating ABA accumulation. Subsequent genetic, physiological, transcriptomic, and molecular investigations revealed that the copper responsive transcription factor SQUAMOSA PROMOTER BINDING PROTEIN-LIKE 7 (SPL7) is a strong regulator of ABA accumulation. We showed that SPL7 is destabilized by high copper and consistently suppresses genes encoding three key oxygenases in the ABA biosynthetic pathway of land plants via binding to the GTAC copper response motifs in their promoters. These results revealed a new mechanism whereby copper availability, inversely reflected by SPL7 abundance, modulates de novo ABA biosynthesis to balance growth and drought tolerance.

**One-sentence summary:** High copper availability represses *SPL7*, releasing its suppression on key ABA biosynthetic genes and leading to increased ABA accumulation that inhibits growth but enhances drought tolerance.

## INTRODUCTION

Due to the autotrophic sessile lifestyle, plants are frequently exposed to environmental conditions that are suboptimal for growth or even threaten their survival. Plants have evolved complex mechanisms to accurately monitor the environment and dynamically adjust metabolism and growth to cope with the abiotic stresses (Claeys and Inzé, 2013; Zhang et al., 2020). Drought is one of the major factors limiting plant habitation and how plants balance growth and drought tolerance has been extensively studied in the past decades (Boyer, 1982; Basu et al., 2016; Gupta et al., 2020). It is now clear that physiological and molecular responses to water limitation are mainly orchestrated by the phytohormone abscisic acid (ABA). Upon dehydration, ABA induce cytosolic calcium oscillations in the guard cells that eventually promote stomatal closure and inhibit stomatal opening to reduce water loss through transpiration (McAinsh et al., 1990; Pei et al., 2000). ABA also programs a substantial portion of the transcriptome, thereby leading to altered metabolism and cellular processes that are required for drought tolerance (Vishwakarma et al., 2017; Chen et al., 2020). In addition, ABA has long been known for its role in active suppression of growth, although the mechanisms are not well understood (Xiong and Zhu, 2003; McAdam and Brodribb, 2018).

The endogenous ABA level is controlled by sophisticated homeostatic mechanisms involving biosynthesis, catabolism, reversible glycosylation, and long-distance transport (Nambara and Marion-Poll, 2005; Umezawa et al., 2010; Vishwakarma et al., 2017; Chen et al., 2020). In land plants, de novo ABA biosynthesis occurs via the carotenoid pathway and involves an array of enzyme-catalyzed steps that initiate in the plastid and conclude in the cytosol (Nambara and Marion-Poll, 2005; Bowman et al., 2017). Zeaxanthin, the first oxygenated carotenoid, is converted to violaxanthin by zeaxanthin epoxidase (ZEP) (Xiong et al., 2002). The 9-cis-epoxycarotenoid dioxygenase (NCED) splits the cis-isomers of violaxanthin and neoxanthin to generate the first cytoplasmic precursor xanthoxin, which is the main rate-limiting step in ABA biosynthesis in Arabidopsis (Iuchi et al., 2001; Endo et al., 2008). *NCED* genes belong to a multigene family and *NCED3* is the major stress-induced isoform in Arabidopsis leaves (Iuchi et al., 2001). The final step is oxidation of abscisic aldehyde to ABA, which is catalyzed by abscisic aldehyde oxidase (AAO). In Arabidopsis, *AAO3* is considered the only *AAO* gene involved in ABA synthesis (Sekimoto et al., 1998; Seo et al., 2000). ABA catabolism proceeds via hydroxylation mediated by members of the cytochrome P450 707A family and conjugation with glucose mediated by glucosyltransferases (Liu et al., 2015). The inactive glucose-conjugated form of ABA can be hydrolyzed by β-glucosidases to replenish cytosolic ABA (Xu et al., 2012). Despite this molecular framework of ABA metabolism, how ABA homeostasis coordinates with other vital physiological processes during growth and development remains to be elucidated.

Copper (Cu) is a trace transition metal in plants that serves as the cofactor for cuproproteins with vital electron transfer or redox reaction functions (Burkhead et al., 2009; Peñarrubia et al., 2015; Zhuang and Li, 2020). In higher plants, Cu is required for growth because the small blue Cu protein plastocyanin, abundantly present in the thylakoid lumen, is an indispensable electron carrier in the Z-scheme of photosynthesis (Molina-Heredia et al., 2003; Weigel et al., 2003). Given that Cu is involved in the synthesis or perception of stress related hormones including ABA (Peñarrubia et al., 2015; Jiang et al., 2021) and ethylene (Rodríguez et al., 1999), is required for the making of protective structures such as lignin (Zhao et al., 2013; Reyt et al., 2020; Zhuang et al., 2020), and participates in various stress responses (Pätsikkä et al., 2002; Abdel-Ghany, 2009; Yamasaki et al., 2009), a role for Cu homeostasis in balancing growth and stress responses warrants investigation.

SPL7 and its orthologs are highly conserved regulators of Cu homeostasis in the green plant lineage, which possess the characteristic Squamosa promoter Binding Protein (SBP) domain that contains the nuclear localization signal and binds the Cu-response DNA motif with a GTAC tetranucleotide core (Kropat et al., 2005; Yamasaki et al., 2009; Zhang et al., 2014). In response to Cu deficiency in Arabidopsis, SPL7 activates genes responsible for increasing cellular Cu uptake (Yamasaki et al., 2009; Bernal et al., 2012; Zhang et al., 2014). SPL7-mediated homeostatic strategy under Cu deficiency also includes activation of the so-called Cu-microRNAs that target transcripts encoding the COPPER/ZINC SUPEROXIDE DISMUTASE (CSD), laccase, and other blue Cu proteins such as plantacyanin (Abdel-Ghany and Pilon, 2008; Yamasaki et al., 2009; Zhang and Li, 2013; Zhao et al., 2015), to ration Cu for immediate usage. In addition to Cu scavenging, SPL7 interacts with other transcriptional regulators, such as ELONGATED HYPOCOTYL 5 that transduces light signaling (Zhang et al., 2014) and CU-DEFICIENCY INDUCED TRANSCRIPTION FACTOR 1 that is required for reproductive development (Yan et al., 2017). These findings suggest that SPL7-mediated Cu homeostasis participates in a broad range of biological processes fundamental to development and responses to the environment.

In this study, we investigated the molecular events underlying an intriguing observation in Arabidopsis that prior Cu treatment retardates growth but enhances draught tolerance at later stages. We found that this balancing act of Cu was exerted by modulating ABA accumulation. Subsequent investigations led to the findings that high Cu destabilizes SPL7, which decreases ABA accumulation by repressing key ABA biosynthetic genes *ZEP*, *NCED3* and *AAO3* via binding to the conserved GTAC Cu-response motifs in their promoters. These findings revealed a potentially conserved mechanism in land plants whereby Cu availability, inversely reflected by SPL7 abundance, modulates de novo ABA biosynthesis via the SPL7-GTAC circuit to balance growth and stress responses.

## RESULTS

### High Cu Hinders Vegetative Growth but Enhances Drought Tolerance

To investigate the role of Cu in balancing growth and stress tolerance in Arabidopsis, we designed a treatment and assaying regime that consists of three plant maintenance phases and three phenotypic checkpoints (Figure 1). In this regime, seed is germinated on agar-solidified half strength MS medium, which contains 0.05 μM Cu (Murashige and Skoog, 1962), with no Cu supplement (referred to as the low Cu condition hereafter) or with Cu added to final concentrations of 10 and 20 μM (referred to as the high Cu condition hereafter). During phase I, seedlings are allowed to grow on the media for up to two weeks before assayed for vegetative growth. In phase II, seedlings are transferred to soil, maintained under well-watered condition, and assayed again for vegetative growth after two weeks. In the final phase, plants are subjected to water withholding for up to three weeks, rewatered for three days, and then assayed for drought tolerance (Figure 1).

**Figure 1.**
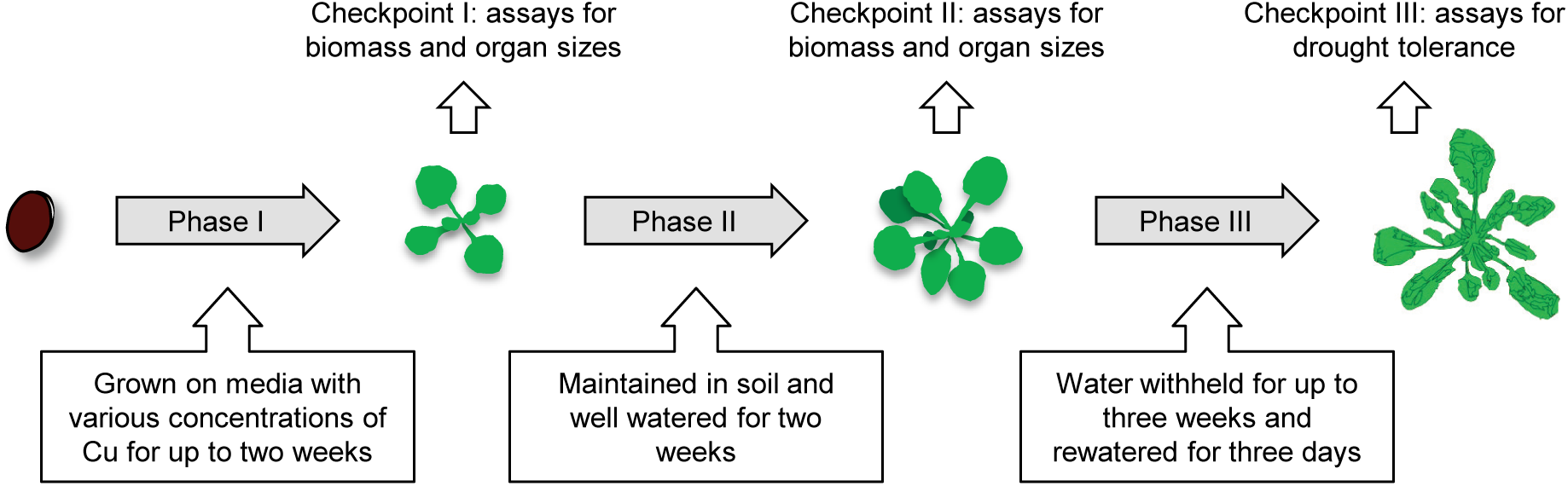
Regime for Cu Treatments and Phenotype Assessments. The regime consists of three plant maintenance phases and three checkpoints for phenotypic assessments. In phase I, seeds are germinated on agar-solidified media supplemented with various concentrations of CuSO_4_ and allowed to grow for up to two weeks. At checkpoint I, seedlings are assayed for vegetive growth including biomass and organ sizes. In phase II, seedlings grown on the media for two weeks are transferred to soil and kept well-watered for two additional weeks. At checkpoint II, the plants are assayed for biomass and organ sizes. In phase III, water is withheld for up to three weeks and re-watered for three days. At checkpoint III, the plants are assayed for drought tolerance.

At checkpoint I, root length of seedlings exposed to high Cu for five days in phase I was already significantly decreased compared to seedlings grown under the low Cu condition (Figure 2). Reduction of foliage area by high Cu was visible after seven days in phase I (Figure 2B). Quantification of 11-d-old seedlings at checkpoint I revealed that high Cu significantly decreased the foliage area and fresh weight based on one-way analysis of variance (ANOVA) (Figure 2C and 2D). Moreover, the reduction of biomass and organ sizes was dependent on the Cu levels as 20 μM Cu supplement resulted in significantly stronger effects on growth than 10 μM Cu (Figure 2D). At checkpoint II, after two-week-old seedlings grown on the media were allowed to recover in soil for two weeks, the impact of high Cu on foliage area and fresh weight persisted (Figure 2E and 2F). These results indicate that high Cu has a lasting effect on inhibiting plant growth and biomass accumulation.

**Figure 2.**
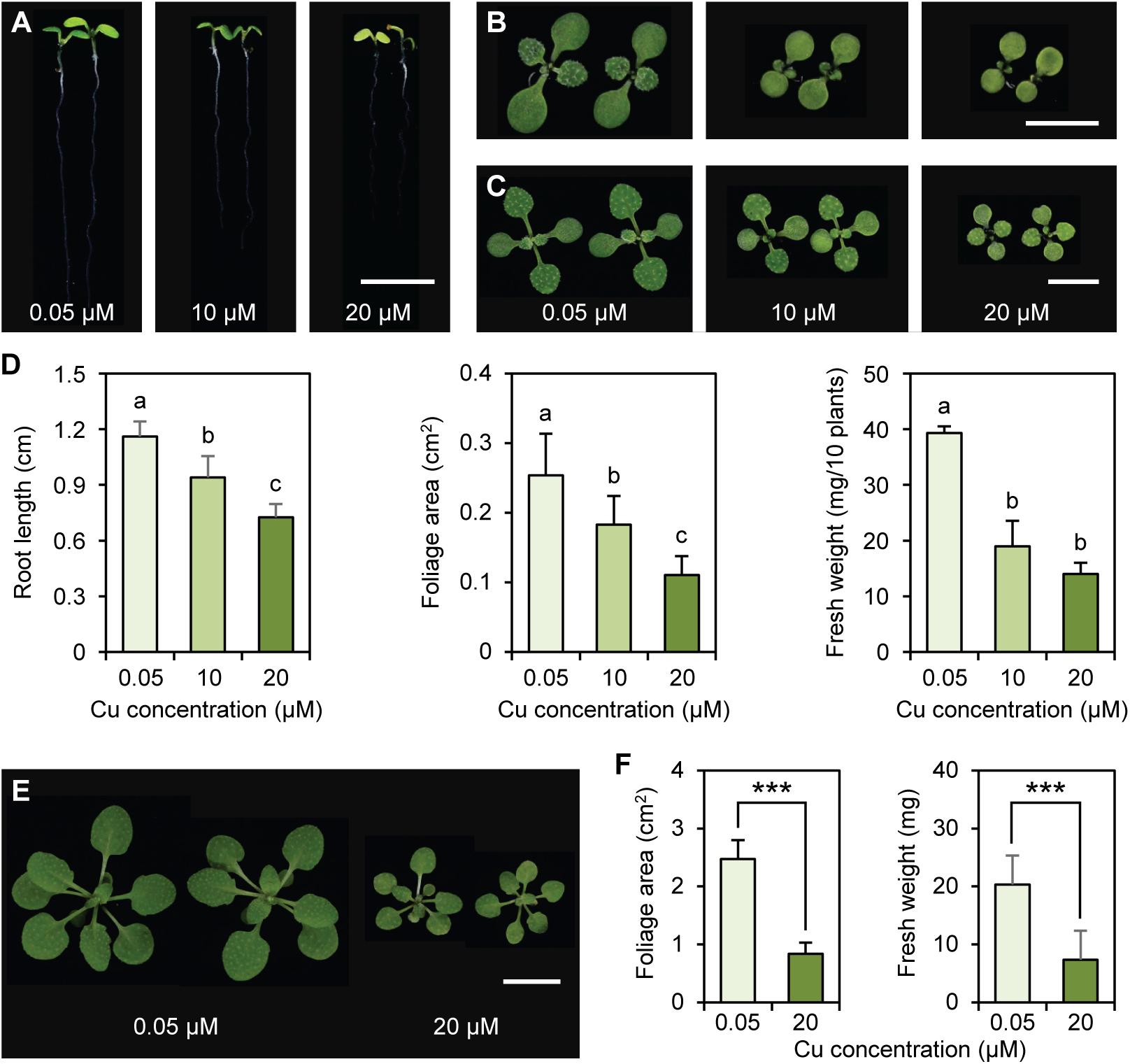
High Cu Hinders Vegetative Growth at Checkpoint I and II. (**A-C**) Representative seedlings at checkpoint I after growing for five (A), seven (B), and 11 days (C) on MS media with the indicated concentrations of CuSO_4_. Bars, 0.5 cm. **(D)** Quantitative measurements of foliage area and fresh weight in 11-d-old seedlings and root length in five-d-old seedlings. Data are means ± SD from n individual plants, where n ≥ 15 for foliage area and root length, and n = 4 for fresh weight (pools of 10 individual seedlings). Different letters denote treatments with significant differences (one-way ANOVA, *p* < 0.01). FW, fresh weight. **(E)** Representative plants at checkpoint II. Seedlings grown on media with the indicated concentration of Cu for two weeks were transferred to soil and photographed two weeks thereafter. Bar, 1 cm. **(F)** Quantitative measurements of foliage area and fresh weight at checkpoint II. Data are means ± SD from n individual plants, where n = 15 for foliage area and n = 4 for fresh weight. ns, not significant; ***, *p* < 0.001 by Student’s *t*-test.

To test the effects of prior Cu exposure on the stress responses, plants in phase III were subjected to water withholding for 18 days and then rewatered for three days (Figure 3). At checkpoint III, we found that, after the water stress was inflicted, the survival rate of plants that had experienced high Cu in the seedling stage was drastically increased in comparison to plants that were exposed to low Cu (Figure 3B). Because plant responses to low water availability include dehydration avoidance and dehydration tolerance (Basu et al., 2016; Gupta et al., 2020; Zhang et al., 2020), we further tested whether both categories are affected by high Cu. Leaf temperature reflects the strength of transpiration with higher temperatures indicating lower levels of transpiration (Wang et al., 2018a). Using infrared thermography, we found that the surface leaf temperature of plants exposed to high Cu was significantly higher than that of the control plants (Figure 3C and 3D). This observation indicates that high Cu exposed plants have a lower transpiration rate and hence reduced water lose than the control plants. We also employed the electrolyte leakage assay to assess cell integrity, which is increasingly compromised during drought stress (Zhao et al., 2016b; Li et al., 2017). We found that the high Cu treated plants displayed significantly reduced membrane permeability than the control plants (Figure 3E), indicating that drought damage was alleviated by high Cu exposure.

**Figure 3.**
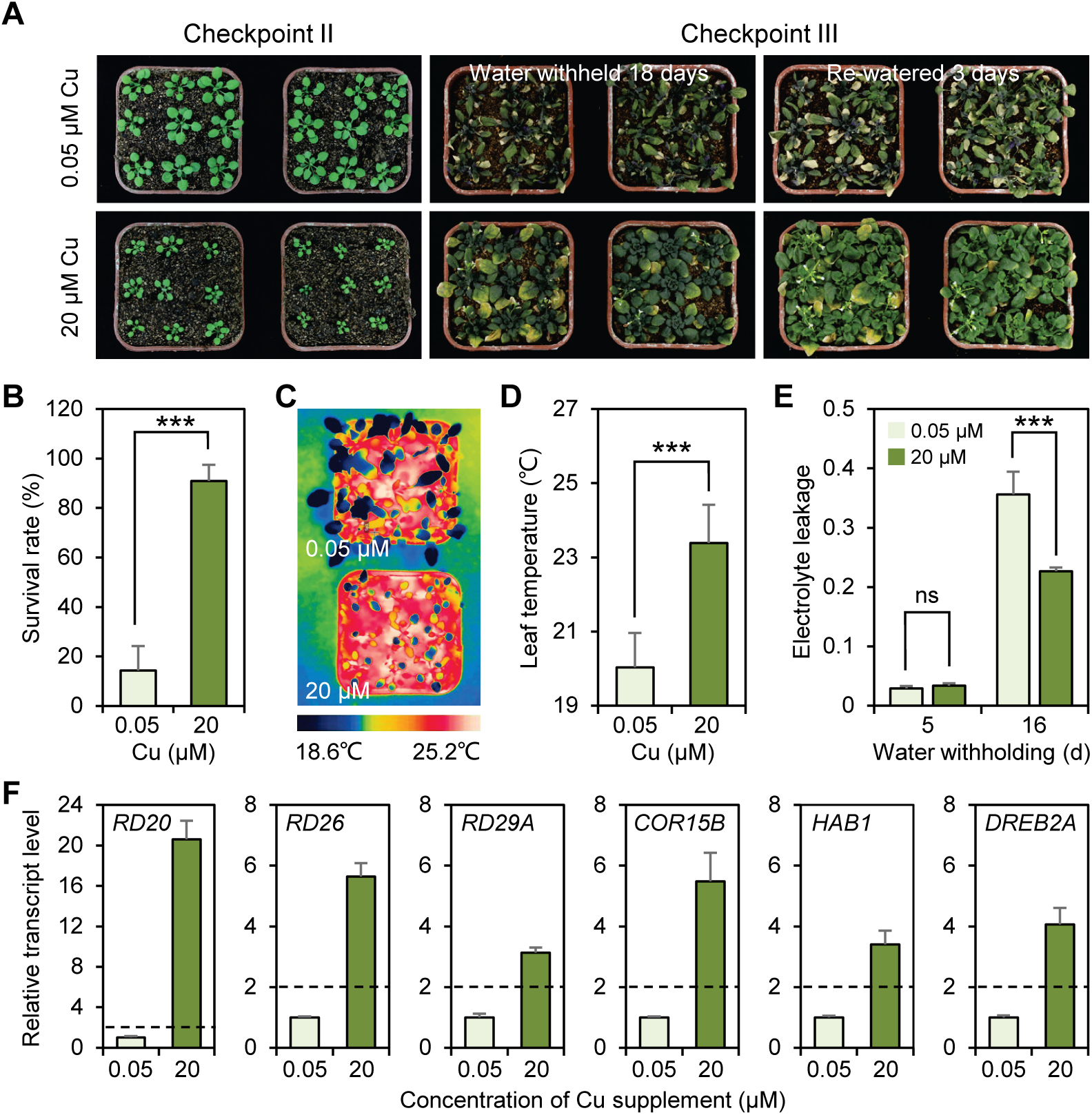
High Cu Enhances Drought Tolerance. **(A)** Photographs of representative pots containing plants at checkpoint II without or with prior high Cu treatment (left), at checkpoint III with water withheld for 18 days (middle), and then re-watered for three days (right). **(B)** Survival rates of plants at checkpoint III after re-watering for three days. The survival rate was determined from six biological replicates each containing nine pots (81 plants). ***, *p* < 0.001 by Student’s *t*-test. **(C)** Representative infrared images of four-week-old plants without or with prior high Cu treatment at checkpoint III. The images were pseudo-colored to show leaf temperature distribution. **(D)** Average leaf temperature quantified from infrared thermography. Data are means ± SD from 20 individual leaves. ***, *p* < 0.001 by Student’s *t*-test. **(E)** Electrolyte leakage at checkpoint III after water withholding for the indicated time. Data are means ± SD from three individual plants. ns, not significant; ***, *p* < 0.001 by Student’s *t*-test. **(F)** Relative transcript abundance of six selected drought-related genes at checkpoint III. Data are means ± SD from three replicate qPCR performed on the same cDNA. Relative transcript abundance was normalized to the wild type and a cutoff of two fold was applied for determining differential expression.

Using reverse transcription coupled quantitative PCR (RT-qPCR), we examined expression pattern of six representative drought-responsive genes, including *RESPONSIVE TO DEHYDRATION 20* (*RD20*), *RD26*, *RD29A*, *COLD REGULATED 15B* (*COR15B*), *HYPERSENSITIVE TO ABA 1* (*HAB1*), *DRE-BINDING PROTEIN 2A* (*DREB2A*) (Umezawa et al., 2010; Chen et al., 2020; Zhang et al., 2020; Zhu et al., 2020). We found that after drought treatment, these hallmark drought-responsive genes were all induced to significantly higher levels (more than twofold) in plants that had experienced high Cu in the seedling stage than in the control plants (Figure 3F). Taken together, these results indicate that prior exposure to high Cu significantly enhances drought tolerance and thus Cu plays an important role in balancing plant growth and drought response.

### SPL7 Is Required for Cu-mediated Growth Regulation

To test whether the Cu responsive transcription factor SPL7 is required for balancing growth and drought tolerance in Arabidopsis, we employed the previously characterized *spl7-1* mutant that contains a T-DNA insertion after the SBP domain (Supplemental Figure 1) (Yamasaki et al., 2009; Zhang et al., 2014). We also generated a deletion mutant using the CRISPR/Cas9 system with paired guide RNAs. The null allele, containing a 4,185 bp deletion that spans the entire coding region, was named *spl7-ko* (Supplemental Figure 1A and 1B). RT-qPCR confirmed that the *spl7-ko* allele had essentially undetectable level of *SPL7* transcript in comparison to the wild type (Supplemental Figure 1C). We also generated the *SPL7_pro_:SPL7 spl7* plants in which the *SPL7* coding region driven by the native *SPL7* promoter was expressed in the *spl7-ko* mutant background. Characterization of the *SPL7_pro_:SPL7 spl7* lines regarding *SPL7* transcript level (Supplemental Figure 1C) and plant morphology (Supplemental Figure 1D) revealed that *SPL7_pro_:SPL7* was able to complement the *spl7* mutation.

The *spl7-1*, *spl7-ko*, and *SPL7_pro_:SPL7 spl7* plants were subjected to the Cu treatment and assessment regime (Figure 1) along with the wild type. Under the low Cu condition, we found that organ sizes (root length and foliage area) and fresh weight of the *spl7-1* and *spl7-ko* mutants were significantly decreased compared to the wild type at both checkpoint I (Figure 4) and checkpoint II (Supplemental Figure 2). The *SPL7_pro_:SPL7 spl7* seedlings, however, exhibited biomass and organ sizes that were statistically undistinguishable from the wild type at both checkpoint I (Figure 4) and checkpoint II (Supplemental Figure 2). These results are consistent with previous reports that SPL7 promotes vegetative growth under low Cu availability (Yamasaki et al., 2009; Bernal et al., 2012; Zhang et al., 2014). When exposed to high Cu during phase I, the wild type and *SPL7_pro_:SPL7 spl7* seedlings exhibited drastically reduced foliage area and fresh weight in comparison to seedlings exposed to low Cu (Supplemental Figure 3). In contrast to the low Cu condition, foliage area and fresh weight of the *spl7-1* and *spl7-ko* seedlings grown under high Cu were statistically undistinguishable from the wild type and *SPL7_pro_:SPL7 spl7* (Supplemental Figure 3). These results indicate that SPL7 is required for manifesting Cu-mediated growth arrest in Arabidopsis.

**Figure 4.**
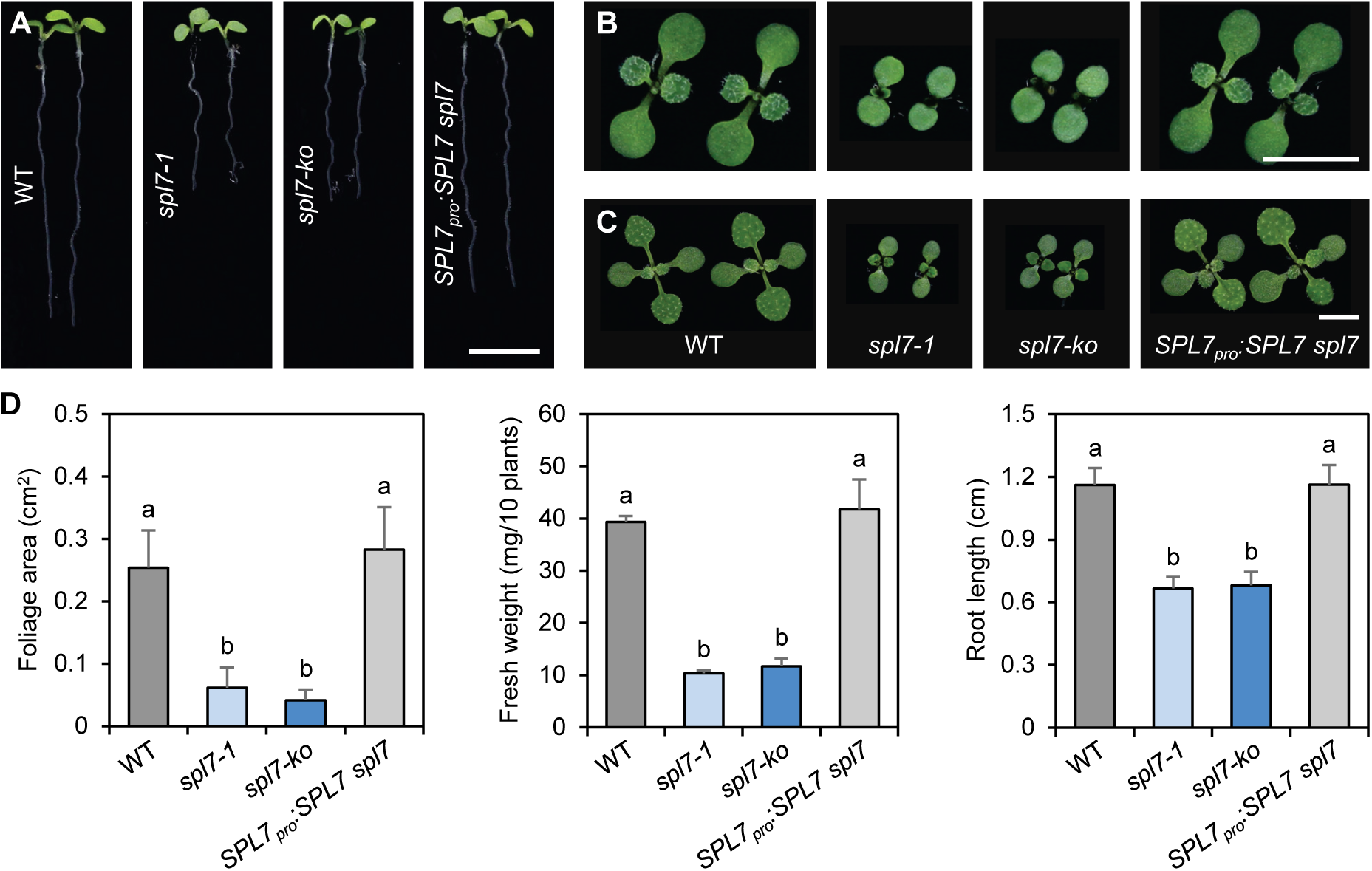
Effects of SPL7 on Vegetative Plant Growth at Checkpoint I. (**A-C**) Representative seedlings of the four indicated genotypes at checkpoint I. Seedlings were grown on the media with 0.05 μM Cu for five (A), seven (B), and 11 days (C) before photographed. Bars, 0.5 cm. (**D**) Quantitative measurements of fresh weight and organ sizes of the four indicated genotypes. Fresh weight and foliage area were measured from 11-d-old seedlings while root length was determined in five-d-old seedlings. Data are means ± SD from n individual seedlings, where n ≥ 15 for foliage area and root length and n = 4 (10 seedlings pooled) for fresh weight. Different letters denote genotypes with significant differences (one-way ANOVA, *p* < 0.001).

### The *spl7* Mutation Enhances Drought Tolerance

To characterize the drought response, the *spl7-1*, *spl7-ko*, and *SPL7_pro_:SPL7 spl7* plants along with the wild type were subjected to water withholding for three weeks during phase III (Figure 5). After this severe drought treatment, we observed that leaves of the wild type and *SPL7_pro_:SPL7 spl7* plants completely withered, but leaves of both *spl7* mutants were dried to a strikingly lesser extent and remained green (Figure 5A). Three days after re-watering, the *spl7* mutants became greener and turgid again while the wild type and *SPL7_pro_:SPL7 spl7* plants failed to recover (Figure 5A). Quantification revealed that the survival rates of the wild type and *SPL7_pro_:SPL7 spl7* plants were both below 5% while the rates were significantly increased to above 95% for both *spl7* mutants (Figure 5B). Examination of the expression profiles of dehydration inducible genes at checkpoint III revealed that, after two weeks of water withdrawing, transcript abundance of *RD20*, *RD26*, *RD29A*, *COR15B*, *HAB1*, and *DREB2A* was significantly increased in the four genotypes (Supplemental Figure 4). Comparison of the degree of induction revealed that five of the six genes (except *DREB2A*) were induced to significantly higher levels (more than twofold) in the *spl7* mutants than in the wild type and *PL7_pro_:SPL7 spl7* plants (Figure 5C). Collectively, these data indicate that the *spl7* mutants display drastically elevated level of drought response.

**Figure 5.**
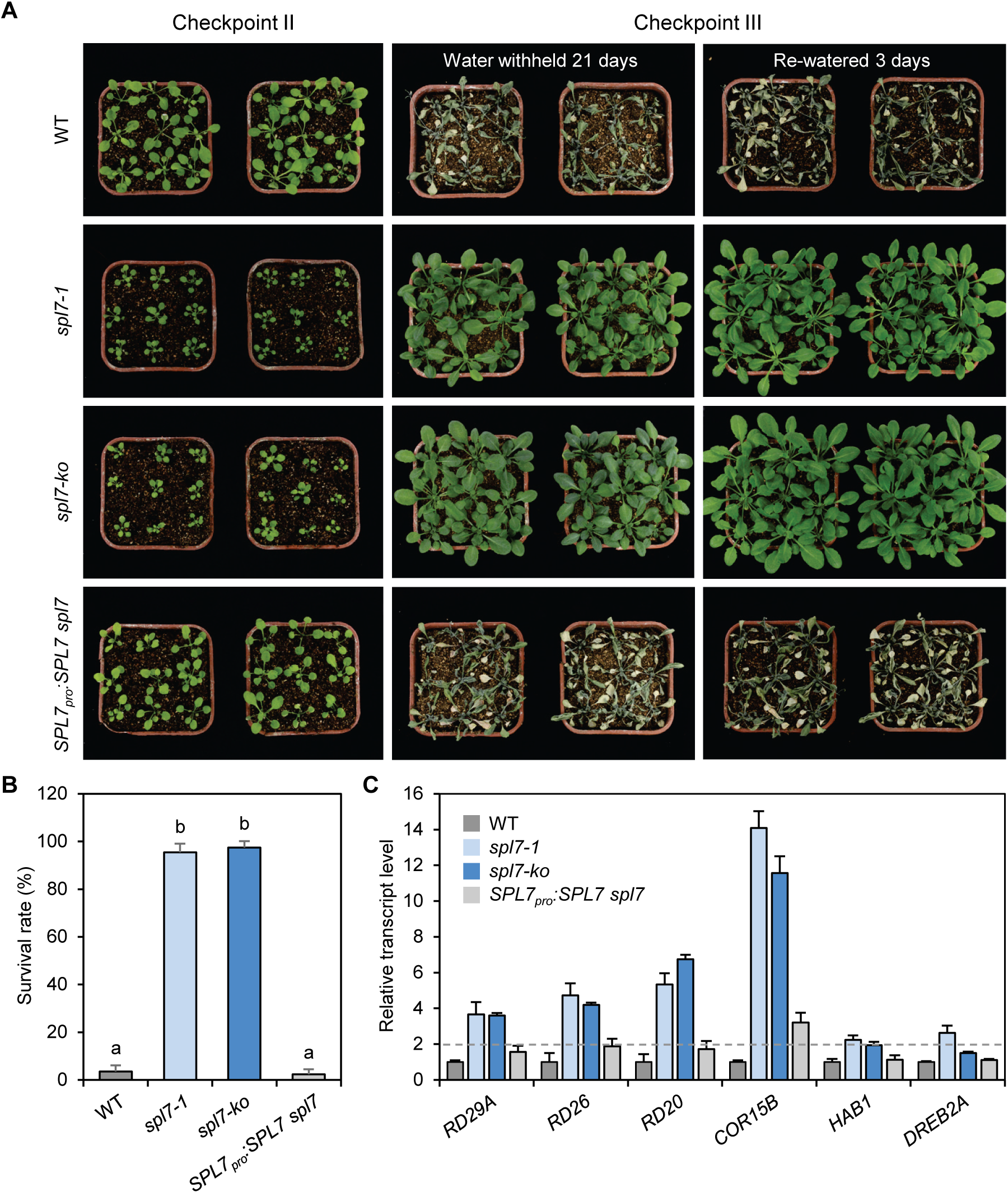
The *spl7* Mutants Are More Resistant to Severe Drought. **(A)** Representative pots containing the indicated genotypes. Photographed from left to right are plants at checkpoint II, checkpoint III with water withheld for 21 days, and then re-watered for three days, respectively. **(B)** Survival rates of plants at checkpoint III after water withholding for 21 days, and then re-watered for three days. Values were determined from five biological replicates each containing nine pots (81 plants). Different letters denote genotypes with significant differences (one-way ANOVA, *p* < 0.001). **(C)** Relative transcript abundance of drought-related genes at checkpoint III. Data are means ± SD from three replicate qPCR performed on the same cDNA. Relative transcript change after and before drought treatment in the four genotypes was normalized to the wild type and a cutoff of two fold was applied for determining differential expression.

We analyzed whether dehydration avoidance and dehydration tolerance were both regulated by *SPL7*. We first examined rosette leaves detached from phase III plants and measured their rates of water loss (Figure 6). We found that both the *spl7-1* and *spl7-ko* leaves showed significantly lower water loss rates than those of the wild type and *SPL7_pro_:SPL7 spl7* (Figure 6A to 6C). We further employed infrared thermography to monitor the surface leaf temperature of the whole plants. This analysis revealed that leaf temperature of the *spl7* mutants was significantly higher than that of the wild type and *SPL7_pro_:SPL7 spl7* plants (Figure 6D and 6E). Together these results indicate that the *spl7* mutants have a lower transpiration rate in the shoots and hence reduced water lose than the wild type plants.

**Figure 6.**
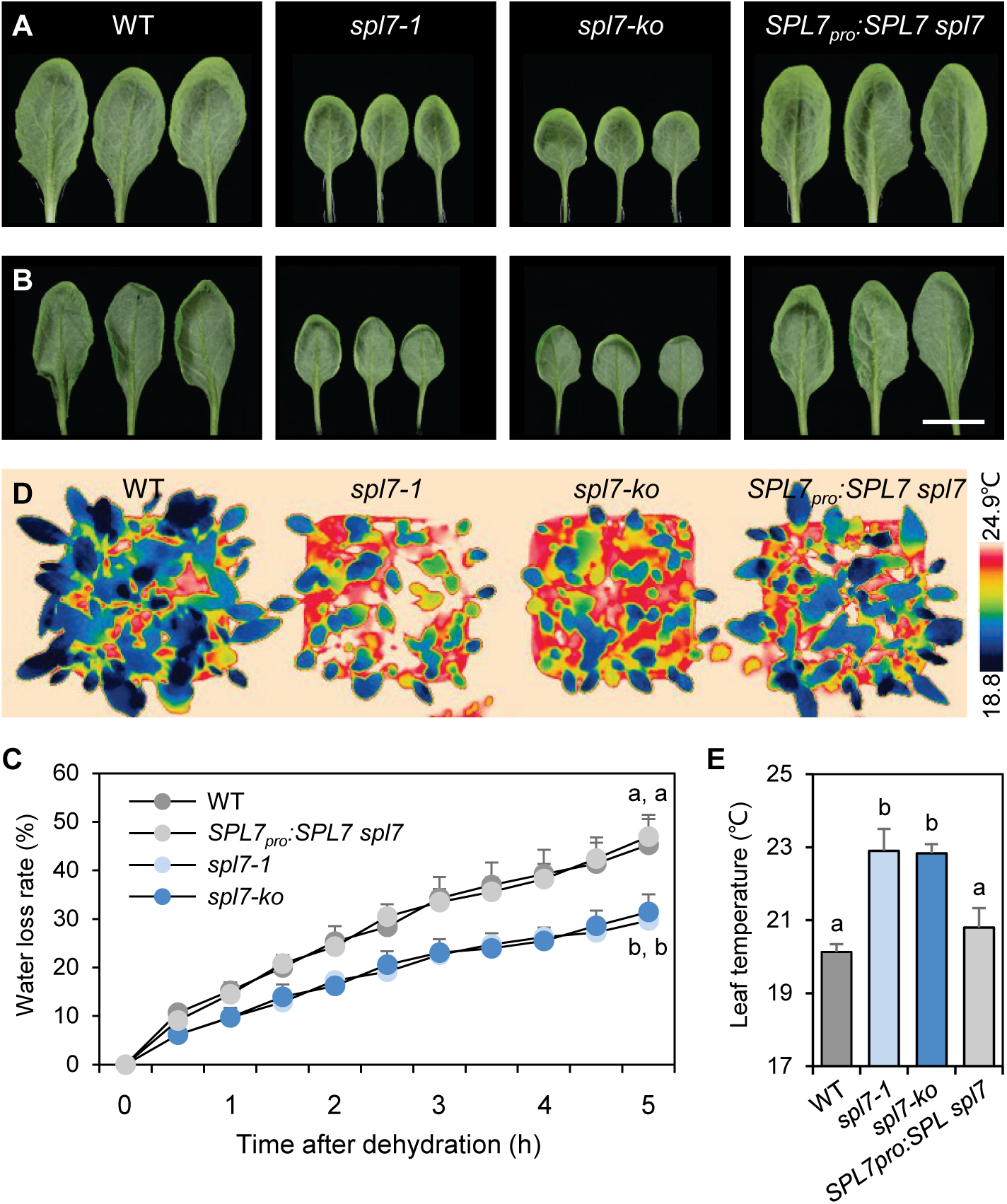
SPL7 Regulates Drought Avoidance. (**A-B**) Representative rosette leaves of the indicated genotypes immediately (A) or three hours after detached from five-week-old phase III plants (B). The leaves were placed in open Petri dishes under room temperature and photographed at the indicate times. Bar, 1 cm. **(C)** Water loss rates of the indicated genotypes. Fresh weight of detached rosette leaves was measured every 30 min over a time course of five hours. Water loss rate was calculated as the proportion of fresh weight lost over the initial weight. Data are means ± SD from three individual experiments each contaning three leaves. Different letters represent genotypes with significant differences at the end of the time course (one-way ANOVA, *p* < 0.001). **(D)** Representative infrared images of four-week-old phase III plants. The images were pseudo-colored to show leaf temperature distribution. **(E)** Quantification of the average leaf temperature from infrared thermography of the indicated genotypes. Data are means ± SD from 20 individual leaves. Different letters denote genotypes with significant differences (one-way ANOVA, *p* < 0.001).

Anthocyanins are known to provide protection to plants under water deprivation (Pourcel et al., 2007; Nakabayashi et al., 2014). We observed that the four genotypes exhibited comparable anthocyanin levels at checkpoint III without water withholding (Figure 7). However, after prolonged water withholding, anthocyanin content increased by more than five folds in the *spl7* mutants in comparison to the wild type and *SPL7_pro_:SPL7 spl7* plants (Figure 7B and 7C). We also used the electrolyte leakage assay to monitor the wild type and *spl7-ko* plants during phase III. We found that by day 15 after water withholding, the *spl7-ko* and wild type plants exhibited the same degree of ion leakage (Figure 7D). However, from day 16 to day 18 after water withholding, the *spl7-ko* plants consistently exhibited significantly lower electrolyte leakage than the wild type (Figure 7D), indicating that membrane damaged by prolonged water shortage was significantly alleviated in *spl7*. Taken together, these results indicate that the *spl7* mutants are more drought-tolerant than the wild type plants.

**Figure 7.**
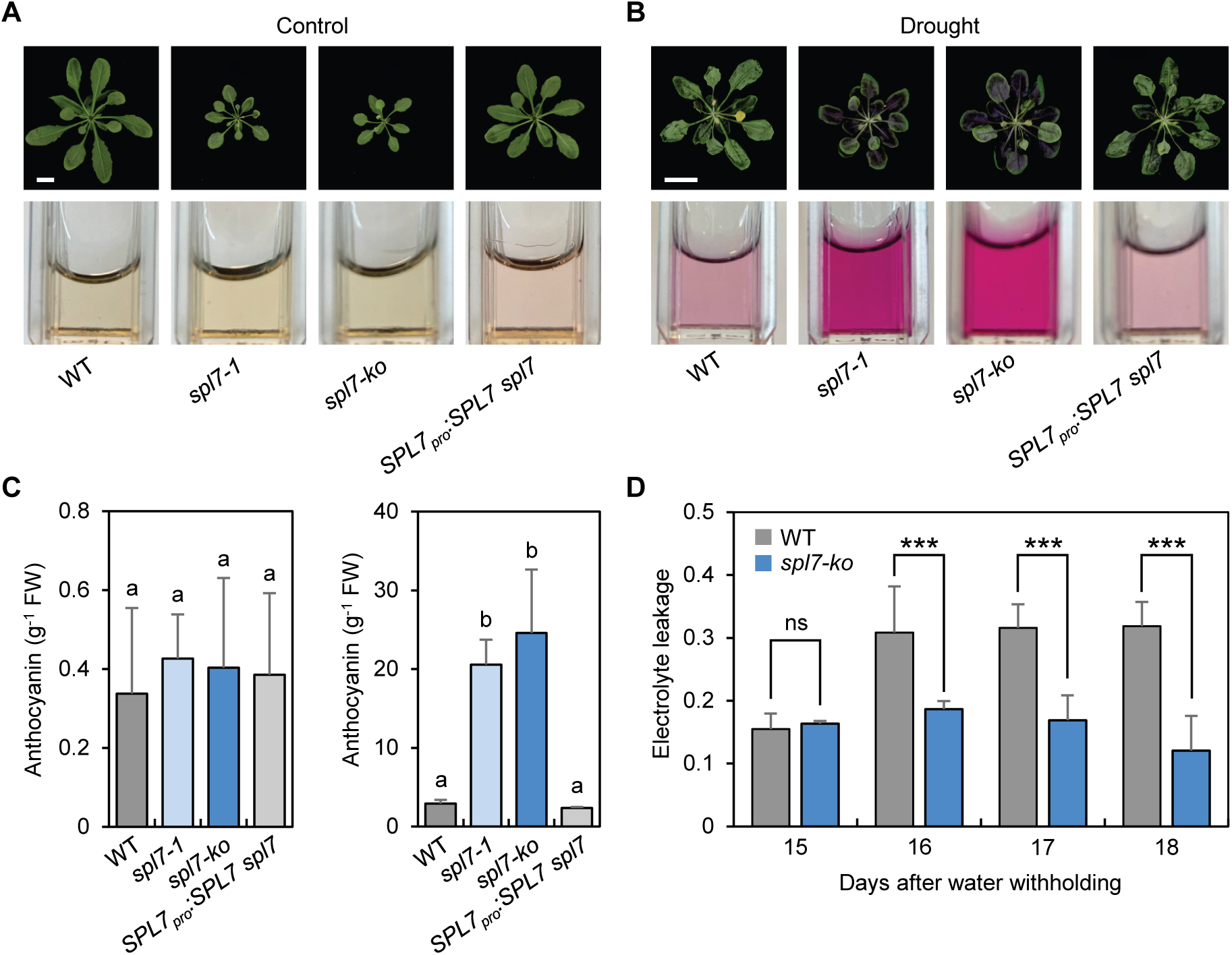
SPL7 Regulates Drought Tolerance. (**A-B**) Phase III plants of the four indicated genotypes were kept well-watered for ten days (A) or subjected to water withholding for ten days (B). Upper panels are representative plants. Bars, 1 cm. Bottom panels are aqueous pigment extractions in cuvettes. **(C)** Quantification of anthocyanin contents in plants without (left) or with drought stress (right). Data are means ± SD from three individual plants. Different letters denote genotypes with significant differences (one-way ANOVA, *p* < 0.001). FW, fresh weight. **(D)** Electrolyte leakage in the wild type and *spl7-ko* plants at checkpoint III after water withholding for the indicated time. Data are means ± SD from three individual plants. ns, not significant; ***, *p* < 0.001 by Student’s *t*-test.

### High Cu Suppresses SPL7 Expression

To elucidate how SPL7 participates in balancing of growth and drought response, we examined *SPL7* expression pattern. We fused the firefly luciferase (LUC) coding region with either the native *SPL7* promoter (*SPL7_pro_:LUC*) or the enhanced cauliflower mosaic virus *35S* promoter (*35S:LUC*). We also fused the SPL7 coding region with that of LUC and placed the chimera protein under control of the native *SPL7* promoter to generate the *SPL7_pro_:SPL7-LUC* reporter. We transformed the wild type Arabidopsis plants and selected transgenic lines expressing the individual reporters (Supplemental Figure 5). The reporter lines were then subjected to the Cu treatment regime shown in Figure 1. As the negative control, we found that luminescence from the *35S:LUC* seedlings at checkpoint I was unaffected by Cu supplementation (Figure 8; Supplemental Figure 6). At checkpoint I, LUC activity in the *SPL7_pro_:LUC* seedlings treated with low and high Cu was not significantly different (Figure 8A; Supplemental Figure 6A). This observation is consistent with previous reports that *SPL7* transcription is not substantially influenced by Cu availability (Yamasaki et al., 2009; Bernal et al., 2012).

**Figure 8.**
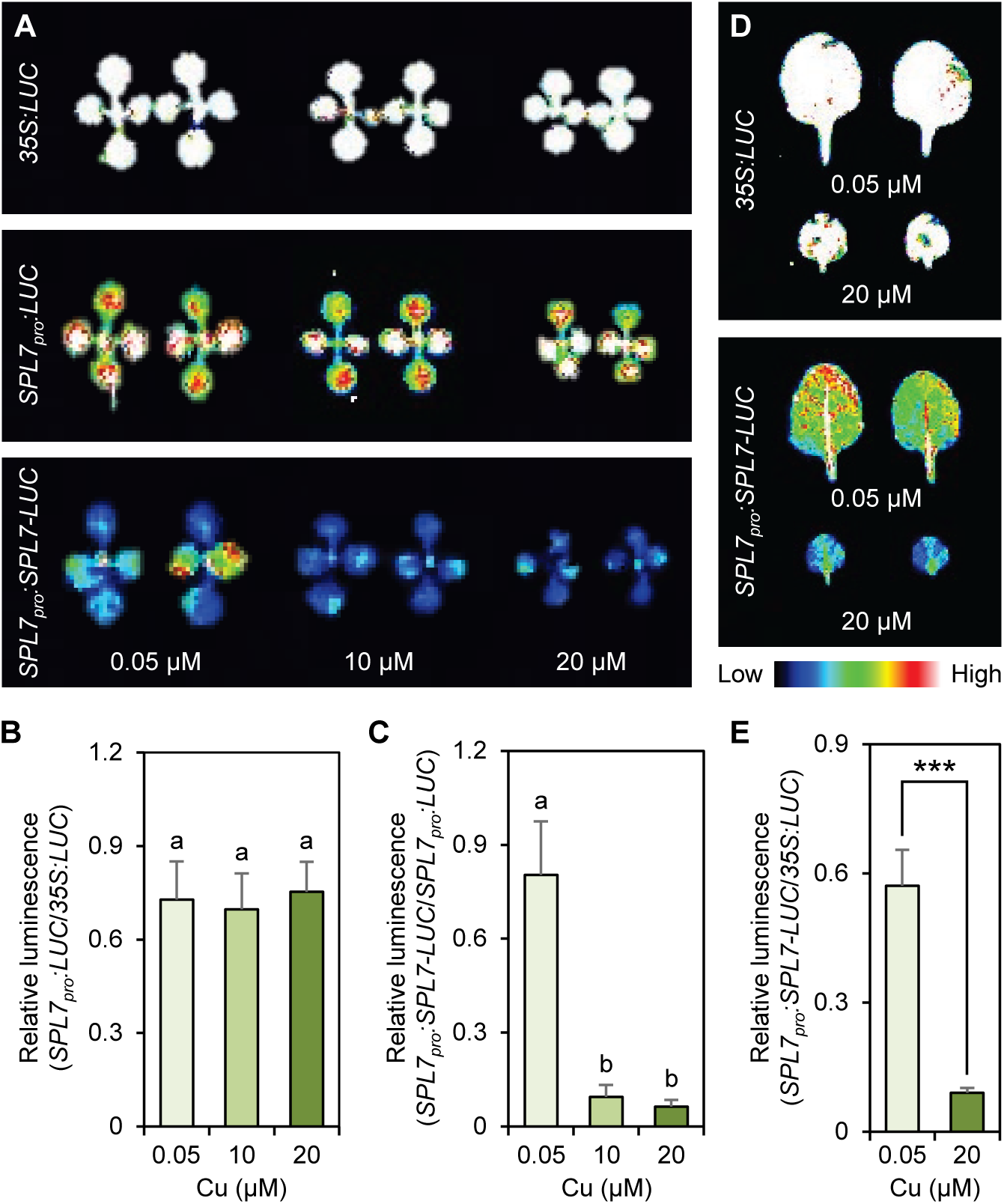
High Cu Reduces SPL7 Abundance. (**A**) Representative luminescence images of the *35S:LUC*, *SPL7_pro_:LUC,* and *SPL7_pro_:SPL7-LUC* seedlings at checkpoint I. Final concentrations of Cu in the media are indicated. (**B-C**) Quantification of relative luminescence from seedlings. LUC luminescence in the *SPL7_pro_:LUC* (B) and *SPL7_pro_:SPL7-LUC* seedlings (C) were normalized to that in the *35S:LUC* and *SPL7_pro_:LUC* seedlings, respectively. Data are mean ± SD from ten individual seedlings. Different letters indicate significant difference (one-way ANOVA, *p* < 0.001). **(D)** Representative luminescence images of detached leaves from the *35S:LUC* and *SPL7_pro_:SPL7-LUC* plants at checkpoint II. **(E)** Quantification of relative luminescence from leaves. LUC luminescence in detached leaves from the *SPL7_pro_:SPL7-LUC* plants were normalized to that in the *35S:LUC* leaves of the same age. Data are means ± SD from ten individual leaves. ***, *p* < 0.001 by Student’s *t*-test.

In contrast, LUC activity at checkpoint I was drastically decreased in the *SPL7_pro_:SPL7-LUC* seedlings experiencing high Cu in comparison to seedlings grown in low Cu (Figure 8A; Supplemental Figure 6A). Normalization of the LUC activity in *SPL7_pro_:LUC* vs. *35S:LUC* (Figure 8B) and in *SPL7_pro_:SPL7-LUC* vs. *SPL7_pro_:LUC* (Figure 8C) revealed that it is the SPL7 abundance that is specifically reduced by high Cu. This finding was confirmed at checkpoint II where we found that LUC activity in *SPL7_pro_:SPL7-LUC* plants with prior exposure to high Cu remained significantly lower than in plants previously exposed to low Cu (Figure 8D and 8E; Supplemental Figure 6B). Therefore, we concluded that high Cu specifically represses SPL7 expression at the protein level, which explains why high Cu and *spl7* mutation reprogram growth and drought tolerance in the same fashion.

### *SPL7*-mediated Cu Homeostasis Modulates ABA Accumulation

To reveal the molecular mechanisms by which SPL7 and Cu balance growth and drought tolerance, we reanalyzed published RNA-sequencing data comparing the wild type and *spl7* plants (Bernal et al., 2012) and the wild type plants treated with or without excess Cu (100 μM). Consistent with previous reports (Bernal et al., 2012), we identified 1155 genes that were differentially expressed between *spl7* and the wild type, of which exactly half (578) were also differentially expressed in response to excess Cu in the wild type (Figure 9; Supplemental Figure 7). Venn diagram (Figure 9A) and clustering analyses (Figure 9B) revealed that 287 of the 578 genes (49.7%) were consistently downregulated while 151 (26.1%) were consistently upregulated in *spl7* and by excess Cu in comparison to the wild type control. Gene Ontology (GO) analysis revealed that the commonly downregulated genes in *spl7* and excess Cu treated wild type plants were significantly enriched with GO terms associated with primary metabolism such as “photosynthesis”, “regulation of photosynthesis”, “carbon fixation”, and “plastid organization” (Figure 9C; Supplemental Figure 7B). This observation explains why growth is inhibited in *spl7* and by high Cu. In contrast, the commonly upregulated genes in *spl7* and by excess Cu were preferentially associated with GO terms related to stress tolerance, such as “response to water deprivation”, “response to salt stress”, “response to oxidative stress”, and “response to abscisic acid” (Figure 9D; Supplemental Figure 7C and 7D). Therefore, both SPL7 and excess Cu globally regulate primary metabolism and response to stresses.

**Figure 9.**
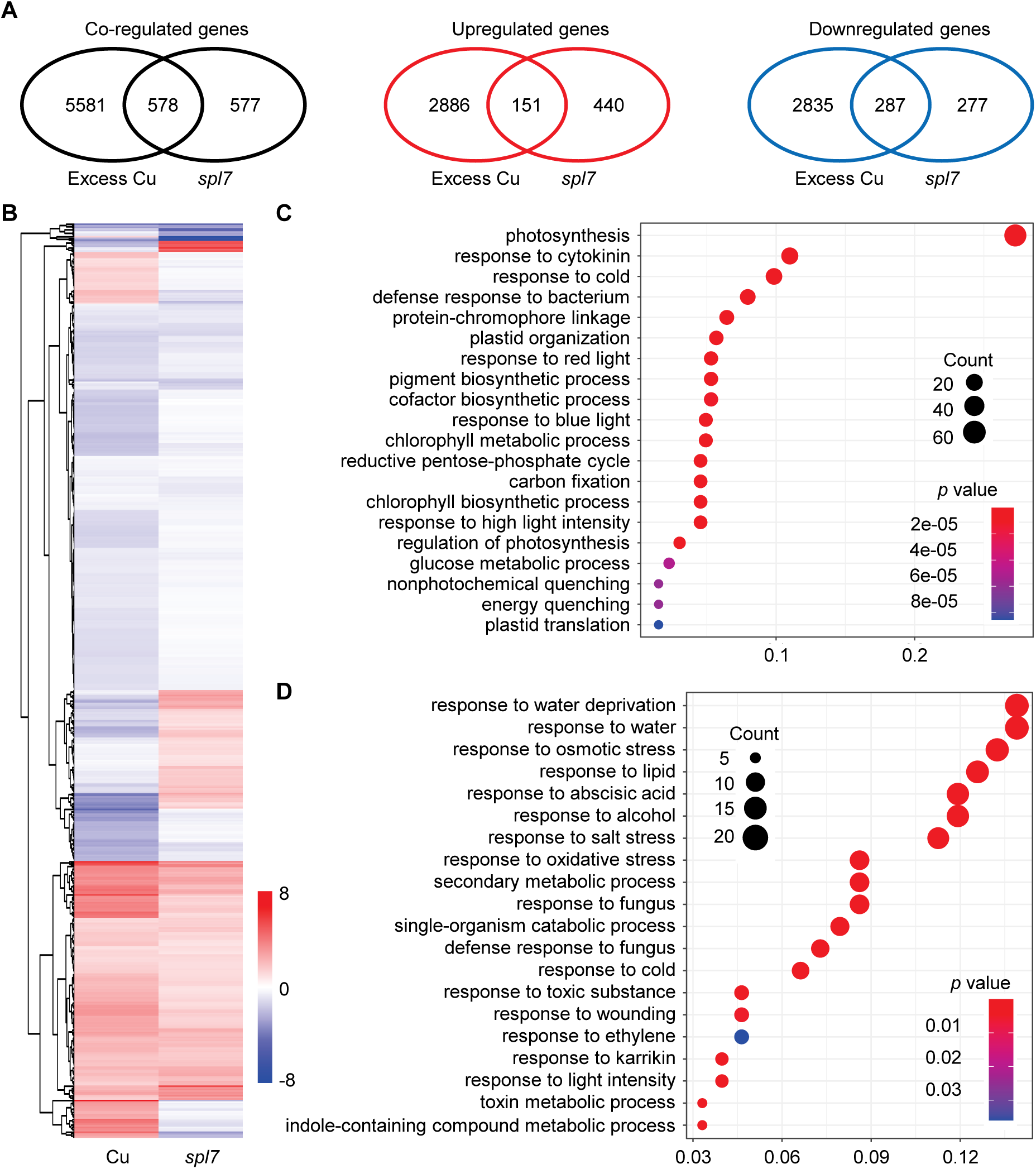
Analysis of Excess Cu and *SPL7* Co-regulated Genes. **(A)** Venn diagram showing the relationship of excess Cu and *SPL7* regulated genes. Differentially expressed genes were identified from RNA-sequencing analysis of excess Cu treated and *spl7* seedlings against the control wild type seedlings. **(B)** Heatmap showing hierarchical clustering of the 578 genes co-regulated by excess Cu and *SPL7*. Colors represent the log_2_-transformed fold changes comparing excess Cu treated seedlings (Cu) and *spl7* seedlings (*spl7*) to the respective controls. (**C-D**) Top 20 enriched GO terms associated with commonly downregulated genes (C) and upregulated genes (D) by excess Cu and in *spl7*. The horizontal axis shows the ratio of the number of co-regulated genes and the number of genes in the genome associated with a given GO term. The color and size of the cycles represent adjusted *p* value and the number of co-regulated genes, respectively.

Given the critical importance of ABA in growth and stress tolerance, we quantitatively compared the levels of endogenous ABA in the *spl7-ko* and wild type plants in our Cu treatment regime. We first measured ABA in seedlings at checkpoint I by the Enzyme-linked immunosorbent assay (ELISA) method using an approved anti-ABA antibody (Ross et al., 1987; Wang et al., 2018b). At checkpoint I, we found that the ABA content in wild type seedlings was significantly increased by Cu treatment (Figure 10). In seedlings exposed to 20 μM Cu, the level of ABA was eight times that of the control seedlings without extra Cu (Figure 10A). In *spl7-ko* seedlings without high Cu exposure, the ABA level was already eight folds of that in the wild type (Figure 10A). ABA level in *spl7-ko* was not further increased by high Cu exposure (Figure 10A). Moreover, we measured ABA content in the wild type and *spl7-ko* plants at checkpoint III using both the ELISA method and the ultrahigh-performance liquid chromatography-triple quadrupole mass spectrometry (UPLC-MS/MS) method (Fu et al., 2012; Jiang et al., 2021). These analyses confirmed that the amount of ABA was significantly higher in *spl7-ko* than in the wild type plants (Figure 10B and 10C). Consistent with the enhanced drought response phenotypes, these results indicate that both high Cu and mutation of *SPL7* substantially increase ABA accumulation.

**Figure 10.**
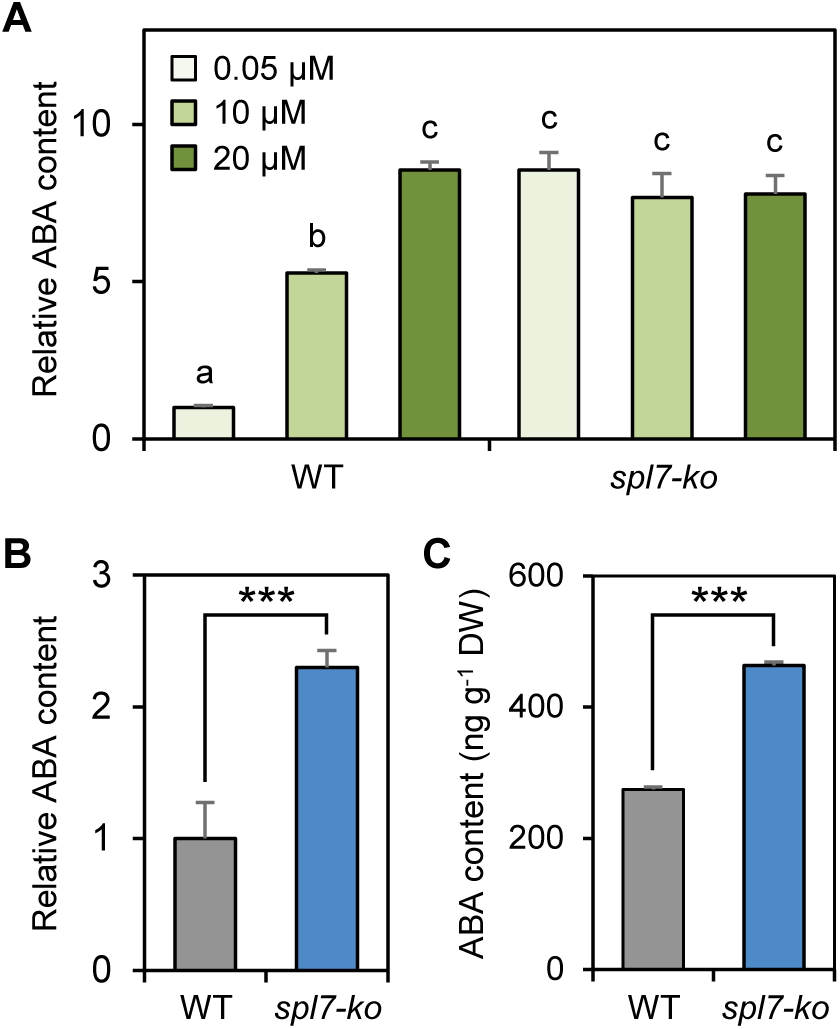
Cu and SPL7 Regulate ABA Accumulation. **(A)** Quantification of the relative levels of endogenous ABA in the wild type and *spl7-ko* seedlings at checkpoint I by the ELISA method. Wild type seedlings grown on the medium with 0.05 μM Cu was used for normalization. Data are mean ± SD from three independent experiments each containing four technical repeats using pooled seedlings. Different letters denote treatments with significant differences (one-way ANOVA, *p* < 0.001). **(B)** Quantification of the relative ABA levels in *spl7-ko* and wild type leaves at checkpoint III by the ELISA method. Data are mean ± SD from three independent experiments each containing four technical repeats using pooled leaves. ***, *p* < 0.001 by Student’s *t*-test. **(C)** Measurement of the absolute ABA levels in *spl7-ko* and wild type leaves at checkpoint III by the UPLC-MS/MS method. Data are mean ± SD from three individual experiments using pooled leaves. ***, *p* < 0.001 by Student’s *t*-test. DW, dry weight.

### SPL7 Directly Suppresses Expression of Key ABA Biosynthetic Genes

To elucidate the mechanism by which SPL7 represses ABA accumulation, we collected 20 structural genes that have been implicated in ABA homeostasis in Arabidopsis (Figure 11; Supplemental Figure 8). Using chromatin immunoprecipitation (ChIP) coupled with whole genome sequencing, we have previously identified 1535 specific SPL7 binding peaks that were associated to 1213 protein-coding genes in Arabidopsis (Zhang et al., 2014). Comparison of this list with the 20 ABA homeostasis genes revealed an overlapping of three genes, namely, *ZEP*, *NCED3*, and *AAO3* (Figure 11B). SPL7 is known to recognize the GTAC core Cu-response DNA motifs that often locate in proximity and form a cluster of several motifs (Yamasaki et al., 2009; Zhang et al., 2014). We found that the proximal promoters of *ZEP*, *AAO3*, and *NCED3* indeed all contain a GTAC cluster (defined herein as neighboring GTAC motifs separated by less than 100 bp) (Figure 11C). To test SPL7 binding to these motifs, we generated the *35S:SPL7^SBP^-GFP* construct in which the *35S* promoter drives the SBP domain of SPL7 fused to the green fluorescent protein (GFP). Using transgenic plants expressing *35S:SPL7^SBP^-GFP*, we performed ChIP with an anti-GFP antibody followed with qPCR analysis. For all three promoters, we found that SPL7^SBP^ occupancy at the GTAC clusters was significantly enriched in *35S:SPL7^SBP^-GFP* relative to the wild type plants, but not in the nearby promoter regions (Figure 11D). These results confirmed that SPL7 directly binds to the promoters of key ABA biosynthetic genes.

**Figure 11.**
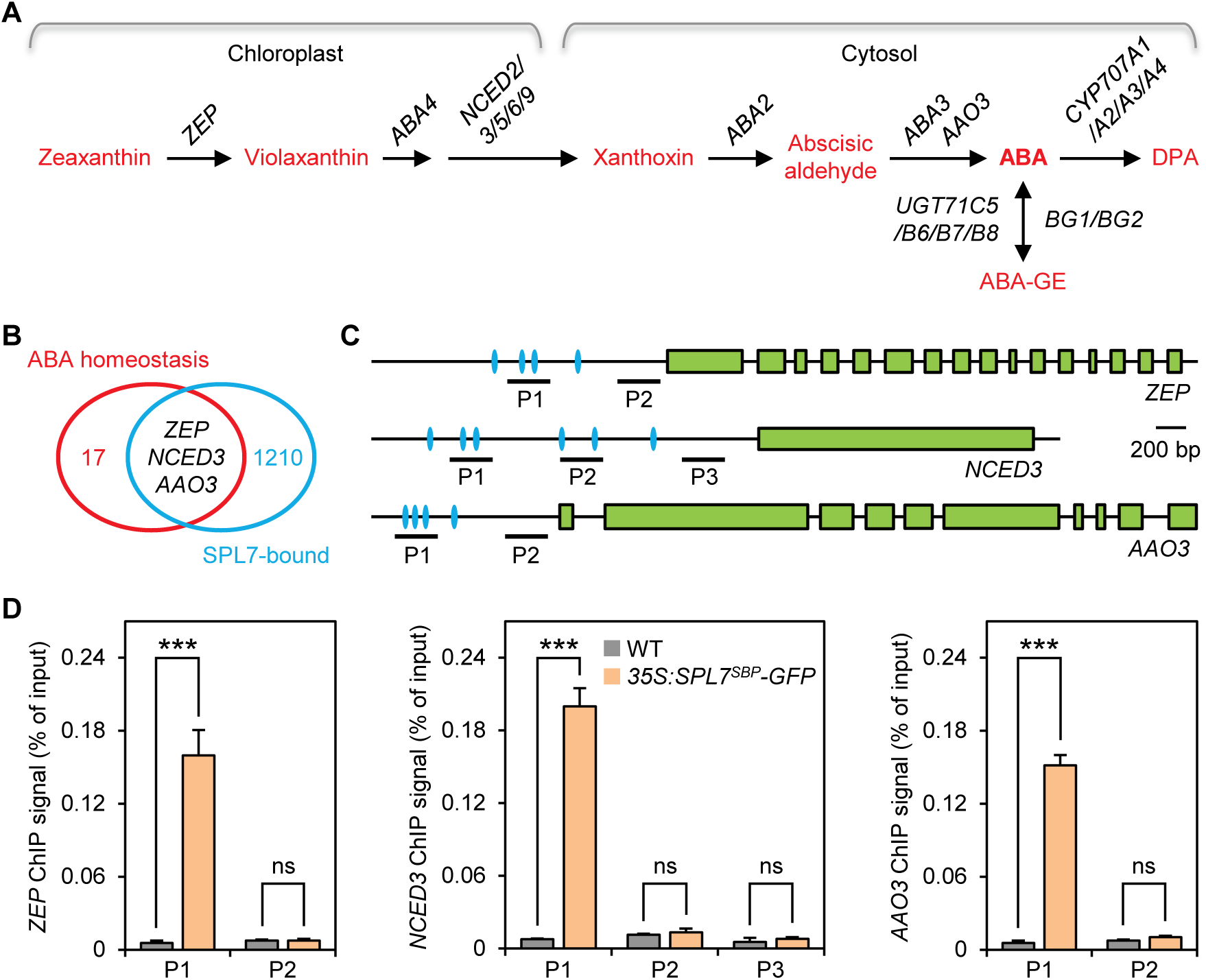
SPL7 Directly Binds to the *ZEP*, *NCED3*, and *AAO3* Promoters. **(A)** Diagram of the simplified ABA homeostasis system in Arabidopsis with genes shown in black and ABA metabolites and catabolites shown in red. ABA-GE, ABA glucosyl ester; DPA, dihydrophaseic acid. *CYP*, *CYTOCHROME P450*; *UGT*, *UDP-GLUCOSYL TRANSFERASE*; *BG*, *BETA-1,3-GLUCANASE*. **(B)** Venn diagram showing overlapping between the 20 ABA homeostasis related genes and 1213 SPL7 associated genes deduced from whole genome ChIP sequencing analysis. **(C)** Schematic drawing of the *ZEP*, *NCED3*, and *AAO3* gene structures. The blue ovals mark positions of the GTAC tetranucleotide in the promoters. Positions of the amplicons subjected to qPCR analysis are marked by horizontal lines. **(D)** Confirmation of SPL7 binding to the *ZEP*, *NCED3*, and *AAO3* promoters by ChIP-qPCR analysis. ChIP was performed using light-grown wild type and *35S:SPL7-GFP* seedlings with or without the anti-GFP antibody. Values are normalized to the respective DNA inputs. Data are ± SD from three qPCR performed on the same DNA. Amplicons are as indicated in A. ns, not significant; ***, *p* < 0.001 by Student’s *t*-test.

To ascertain the effects of SPL7 on the *ZEP*, *NCED3* and *AAO3* promoters, we performed transient assays in tobacco (*Nicotiana benthamiana*) leaves using the LUC and *Renilla* luciferase (REN) dual reporter system (Liu et al., 2014). We generated the *ZEP_pro_:LUC*, *NCED3_pro_:LUC*, and *AAO3_pro_:LUC* reporter constructs by fusing the native *ZEP*, *NCED3*, and *AAO3* promoters with the LUC coding region (Figure 12). We also generated the *miR408_pro_:LUC* and *ACTIN_pro_:LUC* reporter constructs as the positive and negative control, respectively (Figure 12A). For effector constructs, we generated *35S:GFP*, *35S:SPL7-GFP* (in which the *35S* promoter drives the expression the SPL7 coding region fused to GFP), and *35S:SPL7^SBP^-GFP* (Figure 12A). Following co-transfection of tobacco leaf epidermal cells, we found that SPL7-GFP or SPL7^SBP^-GFP, but not GFP alone, significantly increased the LUC/REN ratio of the *miR408_pro_:LUC* reporter (Figure 12B and 12C). This result is consistent with previously report that SPL7 positively regulates the *miR408* promoter (Zhang et al., 2014). As expected, SPL7-GFP and SPL7^SBP^-GFP did not significantly change the LUC/REN ratio of *ACTIN_pro_:LUC* (Figure 12B and 12C). Interestingly, co-expression of SPL7-GFP and SPL7^SBP^-GFP with *ZEP_pro_:LUC*, *NCED3_pro_:LUC*, and *AAO3_pro_:LUC* all significantly reduced the LUC/REN ratio (Figure 12B and 12C), indicating that SPL7 consistently represses the promoters of the three ABA biosynthetic genes.

**Figure 12.**
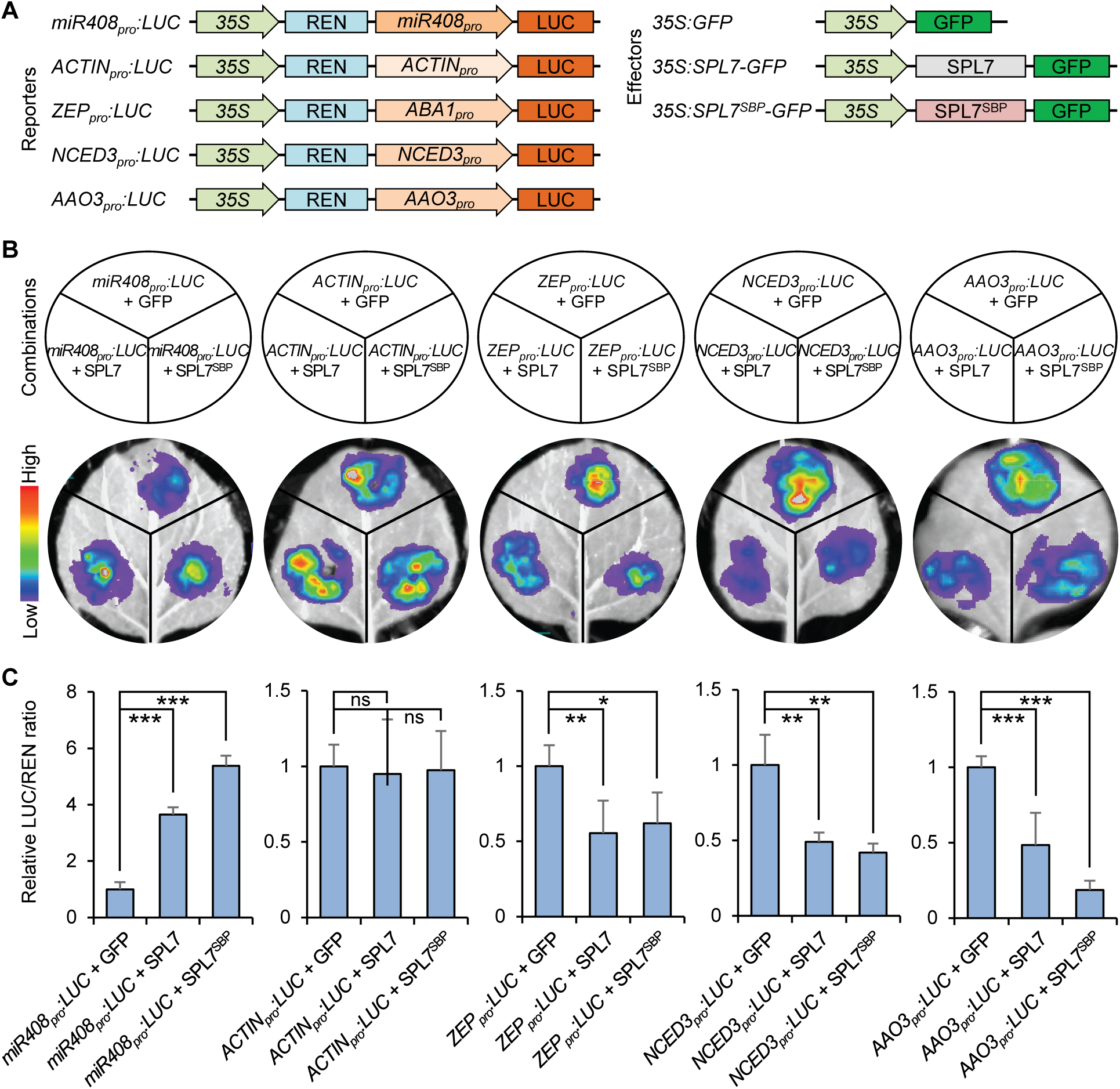
SPL7 Inhibits the *ZEP*, *NCED3*, and *AAO3* Promoters. **(A)** Schematic drawing depicting constructs used for the dual luciferase assay. Constructs containing *LUC* driven by the five tested promoters and *REN* driven by the *35S* promoter (left) were individually combined with one of the three effector constructs (right). The *miR408* and *ACTIN* promoters were used as positive and negative controls, respectively. **(B)** Representative images showing the effects of SPL7 and SPL7^SBP^ on LUC activity driven by the five tested promoters. The indicated combinations of reporter and effector constructs (top) were used to infiltrate tobacco leaf epidermal cells and imaged for LUC activity (bottom). **(C)** Quantification of the relative LUC/REN ratio. Values are means ± SD from three independent transfection events for each combination. ns, not significant; *, *p* < 0.05; **, *p* < 0.01; ***, *p* < 0.001 by Student’s *t-*test.

In the RNA-sequencing data comparing excess Cu treated and *spl7* seedlings against the control, only *NCED3* and *AAO3* were significantly and consistently upregulated among the 20 ABA homeostasis genes (Supplemental Figure 8A). We further found by RT-qPCR analysis that transcript levels of *NCED3* and *AAO3*, but not *ZEP*, in the wild type seedlings were significantly upregulated in high Cu exposed seedlings in comparison with the control seedlings at checkpoint I in our Cu treatment regime (Supplemental Figure 8B). In *spl7-ko*, the expression of *NCED3* and *AAO3* was significantly upregulated without exposure to high Cu while exposure to high Cu did not further increase their expression (Supplemental Figure 8C). Thus, while SPL7 specifically represses the *ZEP*, *NCED3*, and *AAO3* promoters, it substantially influences the expression levels of the latter two in Arabidopsis. These results are consistence with previous reports that *NCED3* and *AAO3* encode the rate-limiting enzymes in ABA biosynthesis and that their differential expression is linked to substantial change in ABA homeostasis (Seo et al., 2000; Iuchi et al., 2001; Chen et al., 2020).

### Suppression of ABA Biosynthetic Genes by *SPL7* Is Conserved in Land Plants

Genomic and genetic analyses have shown that de novo ABA biosynthesis via the carotenoid pathway has arisen and is functionally conserved in land plants (Hauser et al., 2011; Takezawa et al., 2011; Takezawa et al., 2015; Bowman et al., 2017; Komatsu et al., 2020). Furthermore, recognition of the GTAC Cu-response DNA motif by SPL7 orthologs is conserved between the single-cell green alga Chlamydomonas (*Chlamydomonas reinhardtii*) and land plants (Kropat et al., 2005; Yamasaki et al., 2009). These facts prompted us to test whether SPL7-mediated suppression of ABA biosynthetic genes via the GTAC motifs is a conserved feature in land plants. We performed phylogenetic analysis of 12 representative species, including Chlamydomonas, the charophycean green alga *Klebsormidium flaccidum*, the liverwort *Marchantia polymorpha*, the moss *Physcomitrella patens*, the lycophyte *Selaginella moellendorffii*, the gymnosperm *Ginkgo biloba*, the basal angiosperm *Amborella trichopoda*, eudicots Arabidopsis and tomato (*Solanum lycopersicum*), and monocots rice (*Oryza sativa*), maize (*Zea mays*), and the seegrass *Zostera marina*. Consistent with previously reported evolutionary history of the ABA biosynthesis pathway (Bowman et al., 2017), we found that the ZEP and AAO clads include members from the algae while the NCED clade only contains member from land plants (Supplemental Figures 9 to 11). We extracted the 3000 bp proximal promoter regions and searched for the GATC motifs. In the 10 examined land plants, except the *ZEP* in *S. moellendorffii*, we identified at least one member of the *ZEP*, *NCED* and *AAO* clades that contains at least one GTAC cluster in the promoter (Supplemental Figures 9 to 11).

To test whether the GCAT motifs are recognized by SPL7, we selected one representative member of the *ZEP*, *NCED* and *AAO* clades from tomato, rice, and moss (Figure 13; Supplemental Figures 9 to 11). We cloned the nine promoters and generated the *Sly-ZEP_pro_:LUC*, *Osa-ZEP_pro_:LUC*, *Ppa-ZEP_pro_:LUC*, *Sly-NCED_pro_:LUC*, *Osa-NCED_pro_:LUC*, *Ppa-NCED_pro_:LUC*, *Sly-AAO_pro_:LUC*, *Osa-AAO_pro_:LUC*, and *Ppa-AAO3_pro_:LUC* reporter constructs based on the LUC/REN dual luciferase assay in tobacco leaf epidermal cells (Figure 13A). Like the Arabidopsis *ZEP*, *NCED3*, and *AAO3* promoters (Figure 12), we found that all nine tested reporters exhibited significantly decreased LUC/REN ratio in the presence of SPL7^SBP^-GFP, but not GFP (Figure 13B). These results indicate that SPL7 consistently represses the promoters of the ABA biosynthesis genes in both bryophytes and flowering plants. Taken all results together, we concluded that SPL7 ortholog-mediated suppression of key ABA biosynthetic genes is possibly a conserved molecular mechanism for Cu homeostasis to balance growth and drought tolerance in land plants (Figure 14).

**Figure 13.**
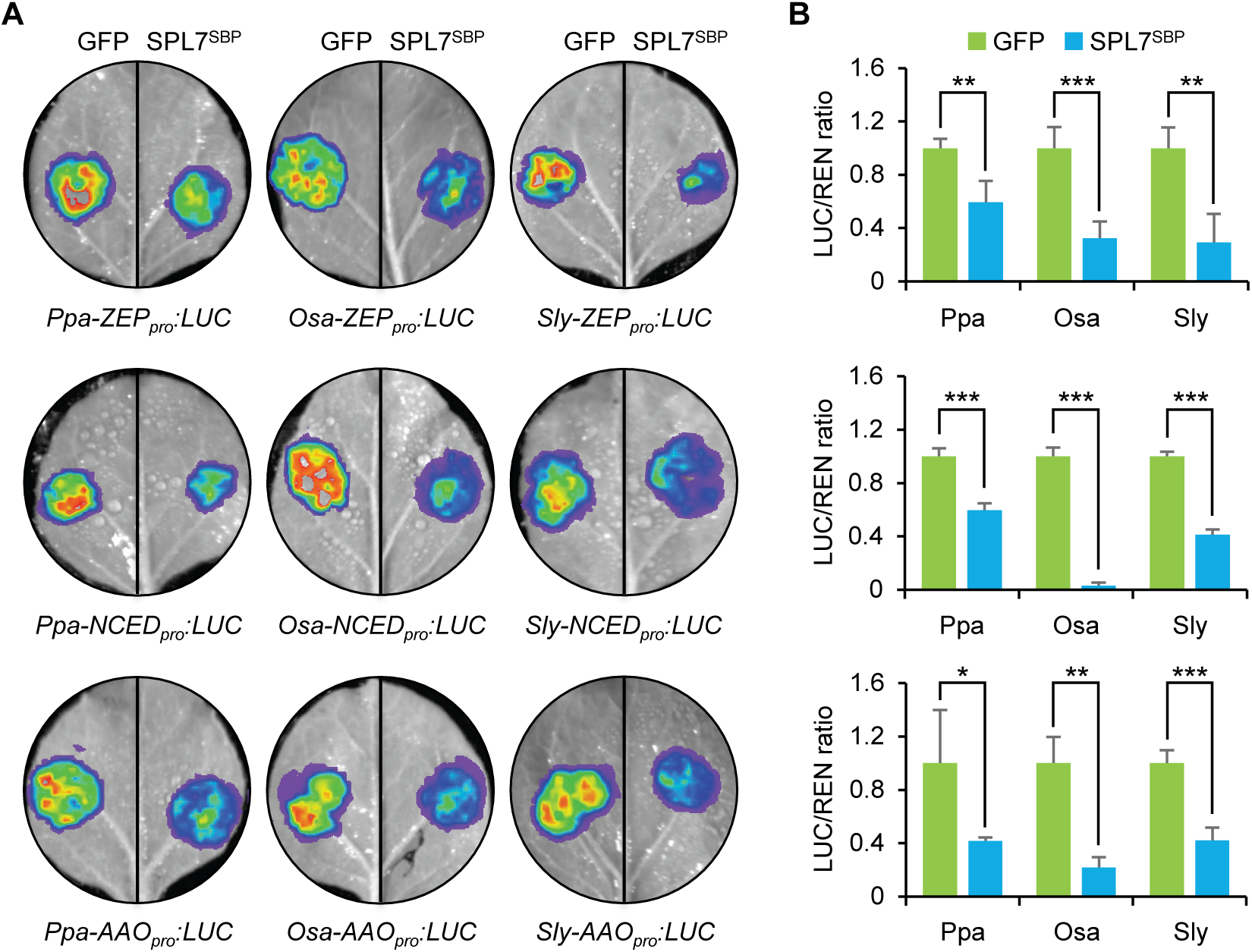
SPL7 Suppresses ABA Biosynthetic Genes in Diverse Land Plants. **(A)** Representative images showing the effects of SPL7^SBP^ on the luciferase activity driven by the tested *ZEP*, *NCED*, and *AAO* promoters selected from moss (Ppa), rice (Osa) and tomato (Sly). Accession numbers of these genes and schematic drawing of their promoters can be found in Supplemental Figures 9-11. **(B)** Quantification of the relative LUC/REN ratio for each tested effector-reporter combination. Values are means ± SD from three independent transfection events. *, *p* < 0.05; **, *p* < 0.01; ***, *p* < 0.001 by Student’s *t-*test.

**Figure 14.**
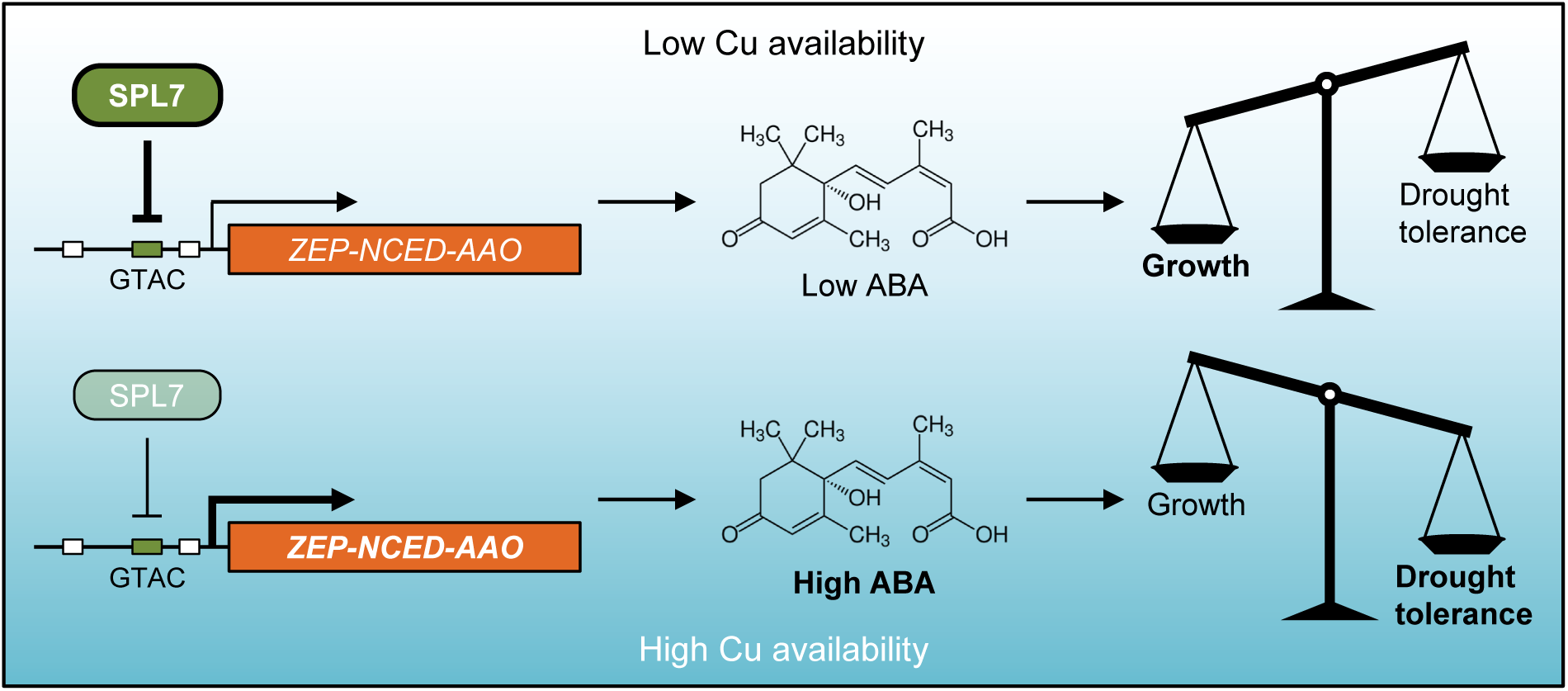
SPL7 Modulates ABA Biosynthesis to Balance Growth and Drought Tolerance. Under low Cu availability, SPL7 represses the *ZEP*, *NCED* and *AAO* promoters through binding to the GTAC Cu-response motifs. This repression mechanism leads to decreased accumulation of cellular ABA, which helps to reserve Cu for growth at the expense of diminished drought tolerance. When Cu supply is in excess, silenced *SPL7* expression leads to de-repression of the key ABA biosynthetic genes, which in turn increases ABA accumulation and enhances drought tolerance. This mechanism is potentially conserved in land plants and SPL7 primarily influences expression of *NCED3* and *AAO3* to modulate ABA biosynthesis in Arabidopsis.

## DISCUSSION

Drought resistance is a complex trait that proceeds through several mechanisms to sense water-deficiency and initiate various coping strategies (Gupta et al., 2020; Zhang et al., 2020). ABA is the main osmostress hormone that regulates molecular and physiological processes, including stomatal closure, root development and architecture, and metabolic reprograming, to promote drought tolerance in plants (Nambara and Marion-Poll, 2005; Umezawa et al., 2010; Vishwakarma et al., 2017; Zhang et al., 2020). However, high ABA is known to actively suppress growth and regulation of ABA level is crucial for maintaining a balance between growth and stress responses (Xiong and Zhu, 2003; Nambara and Marion-Poll, 2005; McAdam and Brodribb, 2018; Zhang et al., 2020). Therefore, elucidating how the ABA homeostasis pathway coordinates with other vital physiological processes to determine cellular ABA content is fundamental to both plant science and agriculture.

### SPL7-mediated Cu Homeostasis Balances Growth and Drought Resistance

Cu, although required only in trace amount, permits life in eukaryotes because of its indispensable functions in electron transfer and redox reactions (Burkhead et al., 2009). The standard MS medium contains 0.1 μM Cu while supplementing the medium with 1 μM Cu was considered optimal for growth of Arabidopsis seedlings (Yamasaki et al., 2009; Garcia-Molina et al., 2014). In this study, we showed that elevated Cu availability (10-20 μM) results in growth retardation (Figure 2) and but allows the plant to perform better under drought (Figure 3). We showed by infrared thermography and electrolyte leakage assays that both dehydration avoidance and dehydration tolerance were enhanced by high Cu (Figure 3). Profiling gene expression and measurements of endogenous ABA further showed that ABA biosynthesis was promoted by high Cu (Figure 9 and 10; Supplemental Figure 7). Collectively, these results demonstrate that increased Cu availability causes the Arabidopsis plant to accumulate more ABA, which restricts growth in the absence of dehydration but increases the ability of the plant to survive severe water deficit in the future.

While the *spl7* mutants grown under low Cu exhibited drastically impeded growth (Figure 4; Supplemental Figure 2 and 3), they showed more restricted water loss of the detached leaves (Figure 6A to 6C), elevated leaf temperature (Figure 6D and 6E), and reduced electrolyte leakage (Figure 7D), which all contribute to increased survival during severe drought (Figure 5). These effects were observed in the wild type plants under high Cu supply (Figure 3). Molecular studies further demonstrated that SPL7 binds to the GTAC motif in the *ZEP*, *NCED3*, and *AAO3* promoters (Figure 11) and consistently represses these promoters (Figure 12). As *ZEP* encodes the enzyme that initiates the ABA pathway, *NCED3* encodes the enzyme that provides the first cytoplasmic intermediate and is the rate-limiting step, and *AAO3* encodes the enzyme that catalyzes the final step (Seo et al., 2000; Iuchi et al., 2001; Nambara and Marion-Poll, 2005), their repression by SPL7 is clearly strategically placed. Accordingly, we found that the level of ABA drastically increased in the *spl7* mutants in comparison to the wild type (Figure 10). These findings indicate that SPL7 is required for manifesting the balancing effect of Cu on growth and drought tolerance.

Consistent with its role as a Cu-responsive transcription factor under Cu deficiency (Kropat et al., 2005; Yamasaki et al., 2009; Bernal et al., 2012; Zhang et al., 2014), we found that SPL7 is silenced under high Cu (Figure 8; Supplemental Figures 5 and 6). These results support a scenario in which SPL7 regulates Cu partitioning based on the prevailing Cu condition to balance growth and drought tolerance (Figure 14). Under low Cu, in addition to activating the Cu-microRNAs that favor Cu allocation to the plastocyanin for photosynthesis (Yamasaki et al., 2009; Zhang et al., 2014; Pan et al., 2018; Jiang et al., 2021), direct repression of key ABA biosynthetic genes by SPL7 results in diminished ABA production to promote growth (Figure 4 and 10 to 12). Under high Cu, degradation of SPL7 releases suppression on ABA biosynthesis and high ABA in turn enhances stress tolerance. Thus, SPL7-mediated Cu homeostasis provides a mechanism linking primary metabolism to de novo ABA production and thereby orchestrating growth and drought resistance.

### Biochemical Bases for Coordinating Cu and ABA Homeostasis

Because Cu is highly reactive, eukaryotes have evolved a strong chelating capacity in the cytosol that is estimated to allow for less than one free Cu ion per cell (Rae et al., 1999; Finney and O’Halloran, 2003). Consequently, elaborate transferring systems are present in eukaryotic cells for the precise delivery of Cu to specific target proteins (Burkhead et al., 2009; Robinson and Winge, 2010). These features place the transition metal in a unique position to balance growth and ABA production in plants. One of the biochemical bases for Cu impacting ABA biosynthesis is AAO acting in the cytosol. In Arabidopsis, *AAO3* encodes the aldehyde oxidase that converts abscisic aldehyde to ABA, the last step of de novo ABA biosynthesis (Seo et al., 2000). AAO3 is a molybdoenzyme that requires the molybdenum cofactor for catalytic activity (Seo et al., 2000; Xiong et al., 2001; Kuper et al., 2004). Structural studies of CO-FACTOR FOR NITRATE REDUCTASE AND XANTHINE DEHYDROGENASE 1 (Cnx1) demonstrated that formation of a Cu-dithiolate complex is a critical step in molybdenum cofactor biosynthesis, which protects the reactive dithiolate before molybdenum insertion (Kuper et al., 2004). It was thought that an unidentified chaperone is required in the cytosol for passing on Cu to the dithiolate group for making the Cu-dithiolate complex and ultimately determining AAO activity (Kuper et al., 2004; Peñarrubia et al., 2015).

CSD1 is a major Cu sink protein in the cytosol that functions to detoxify superoxide radicals (Burkhead et al., 2009). CSD1 is a target for SPL7-mediated scavenging under Cu deficiency as its expression is post-transcriptionally silenced by SPL7-activated miR398 (Abdel-Ghany and Pilon, 2008; Burkhead et al., 2009; Yamasaki et al., 2009). It could be extrapolated that Cnx1, which requires Cu for synthesizing the molybdenum cofactor in the cytosol, is regulated by SPL7-meidated Cu scavenging. In this scenario, elevated Cu availability represses *SPL7* expression to retain more Cu in the cytosol, which in turn elevates AAO3 activity and hence de novo ABA biosynthesis. Although future experiments are needed to prove this mechanism, it is consistent with previous models of cellular Cu partitioning. On the one hand, the level of chloroplastic Cu correlates with the abundance of functional plastocyanin and reflects photosynthetic performance (Burkhead et al., 2009; Zhang et al., 2014; Pan et al., 2018). On the other hand, formation of the molybdenum cofactor, which is required for AAO activity (Seo et al., 2000; Kuper et al., 2004), elevates cellular ABA content. Partitioning of the plastocyanin pool of Cu in the chloroplast and the cytosol pool of Cu would afford the plants a “seesaw” mechanism to adjust growth and stress resistance (Figure 14).

In plants, an important constituent of the cellular Cu homeostasis machinery is the CTR-like high-affinity transporters, denoted COPTs, that translocate Cu ions across the membrane (Puig and Thiele, 2002; Pope et al., 2012). Arabidopsis possesses six COPTs named COPT1-COPT6 that perform specialized functions in Cu homeostasis. In silico analysis of the *COPT* promoters in Arabidopsis revealed that the main ABA responsive cis-elements, namely ABRE (ABA-responsive element), MYB/MYC, and DPBF (Dc3-Promoter Binding Factor), are abundantly present (Peñarrubia et al., 2015), suggesting that these transporters could be regulated by ABA. Thus, osmostress induced changes in Cu uptake and partitioning via COPTs provide another possible biochemical basis for coordinating Cu and ABA homeostasis.

### Cu and ABA, an Ancient Balancing Act in Land Plants?

Protecting cells from dehydration is critical for sessile plants to survive water-limited terrestrial environments. Genetic and genomic analyses have shown that de novo ABA biosynthesis via the carotenoid pathway is likely functionally conserved in land plants, which are considered to have descended from an ancestral form of aquatic algae (Hauser et al., 2011; Takezawa et al., 2011; Takezawa et al., 2015; Bowman et al., 2017; Komatsu et al., 2020). Likewise, genomic and genetic evidence also strongly suggests that the core ABA signaling pathway was already present in the common ancestor of land plants (Khandelwal et al., 2010; Bowman et al., 2017; Eklund et al., 2018; Sun et al., 2019; Komatsu et al., 2020). It was proposed that ABA functions are a major adaptation in early land plants, enabling them to balance protection against osmostress and growth for successful colonization of terrestrial environments (Khandelwal et al., 2010; Takezawa et al., 2011; Eklund et al., 2018; Sun et al., 2019; Komatsu et al., 2020).

We found several lines of evidence suggesting that SPL7-mediated suppression of ABA biosynthetic genes is a conserved feature in land plants. First, both SPL7 orthologs and recognition of the GTAC Cu-response DNA motif by SPL7 orthologs are deeply conserved in the green plant lineage (Kropat et al., 2005; Yamasaki et al., 2009; Zhang et al., 2014). Second, we showed in the current study that functional GTAC Cu-response motifs are present in the promoters of key ABA biosynthetic genes in phylogenetically representative land plant species (Figure 12 and 13; Supplemental Figure 9 to 11). Additionally, the phenomenon that high Cu increases ABA accumulation has been reported by physiological studies in different plant species (An et al., 2008; Ye et al., 2014; Jiang et al., 2021). Repression of ABA biosynthesis by the SPL7-GTAC circuit thus appears to be a conserved mechanism in land plants.

Based on the above considerations, we hypothesize that concurrent increases in cytosolic Cu concentration and ABA biosynthesis upon dehydration may have facilitated molecular evolution in early land plants to couple the two homeostatic events via the SPL7-GTAC circuit, enabling them to better balance desiccation tolerance under water deficiency and rapid growth under water sufficiency. This feature may have given these plants an advantage in terrestrial environments that allowed them to establish a firm foothold on land (Takezawa et al., 2011; Komatsu et al., 2020). Future studies of osmostress responses and Cu homeostasis in basal streptophytes and bryophytes would add to our understanding of land plant evolution. On a practical note, drought is now an increasing problem worldwide that adversely affects plant growth and crop production (Gupta et al., 2020; Zhang et al., 2020). Given the evolutionary conservation of SPL7-mediated coordination of Cu and ABA homeostasis, we anticipate this mechanism to be tapped into for engineering crops with higher efficiency of water usage to alleviate the global drought challenge.

## METHODS

### Plant Materials and Growth Conditions

The wild-type line used in this study was *Arabidopsis thaliana* ecotype Col-0. To produce a complete loss-of-function *spl7* mutant, a pair of sgRNAs were designed to target both ends of the *SPL7* coding region and were inserted into the pAtU6-26-SK vector. The two sgRNA cassettes were digested out of pAtU6-26-SK using the restrict enzymes by *Spe*I and *Nhe*I (New England Biolabs) and subcloned into pCAMBIA1300-35S-Cas9 to generate pCAMBIA1300-AtU6-26-sgRNA1-AtU6-26-sgRNA2-35S-Cas9. The construct was used to transform wild type plants following the standard floral dip method (Clough and Bent, 1998) and selected with 50 mg L^-1^ Hygromycin (Roche). A total of 60 T_1_ plants were individually genotyped by PCR and sequencing. Four lines were selected for propagation of which three were confirmed by genotyping at the T_4_ generation as Cas9-free homozygous *spl7-ko* lines. To complement *spl7-ko*, the native promoter and the full-length coding region of *SPL7* were cloned into the same pCAMBIA1381 vector between the *BamH*I and *Pst*I sites and between the *Spe*I and *BstE*II sites, respectively. The construct was subsequently transformed into the *spl7-ko* plants. Following transformation and selection, Hygromycin resistant transformants were allowed to propagate to the T_3_ generation. Three independent homozygous *SPL7_pro_:SPL7 spl7* lines were selected by genotyping. To produce the *35S:LUC*, *SPL7_pro_:LUC*, and *SPL7_pro_:SPL7-LUC* constructs, the enhanced cauliflower mosaic virus *35S* promoter, native *SPL7* promoter, and the native promoter together with the full-length SPL7 coding region were individually cloned into the p1305.1-LUC vector between the *BamH*I and *Nco*I sites. Following transformation, homozygous transformants were selected by Hygromycin resistance and genotyping in the T_3_ population. Three independent homozygous *35S:LUC*, *SPL7_pro_:LUC*, and *SPL7_pro_:SPL7-LUC* lines were selected for subsequent analyses. To produce the *35S:SPL7^SBP^-GFP* construct, *SPL7^SBP^* was cloned the into pJIM19-GFP vector in frame of the *GFP* coding region. Following transformation and selection, Hygromycin resistant transformants were allowed to propagate to the T_3_ generation. Three independent homozygous *35S:SPL7^SBP^-GFP* lines were selected by genotyping. Primers used for the generation of various constructs are listed in Supplemental Table 1.

To grow Arabidopsis plants, surface sterilized seeds were plated on agar-solidified half strength MS media including 1% (w/v) sucrose and incubated in the dark at 4°C for three days. Germinated seedlings were allowed to grow on plates for up to two weeks in a Percival culture chamber (CU-36L5) under the conditions of 16 h light/8 h dark at 22°C/20°C with full lights on. Seedlings were transferred to commercial soil and maintained in a Percival growth chamber (E-36L) under the settings of 8 h light/16 h dark cycle at 22°C/20°C and 50% relative humidity. Light was provided by white fluorescent bulbs with intensity of approximately 120 μmol m^-2^ s^-1^. Transient assays were performed using tobacco plants, which were grown in a walk-in growth chamber under a 16 h light/8 h dark cycle, 25°C/21°C, 50% relative humidity, and light intensity of approximately 150 μmol m^-2^ s^-1^.

### Drought Tolerance Assays

To monitor water-loss of single detached leaf, rosette leaves of various genotypes at the same developmental stage were excised from five-week-old plants and placed in open Petri dishes under room temperature. Fresh weight of the detached leaves was measured every 30 min until the indicated times. Water loss was calculated as the proportion of total weight lost over the initial fresh weight. Three independent experiments were performed each including three or four plants. For drought treatment of whole plants, seedlings grown on half-strength MS medium supplemented with the indicated concentrations of Cu for two weeks were transferred to the soil. Nine seedlings were placed in each pot. Care was taken to put the same amount of soil in each pot and the same volume of water in each flat (occupied by 18 pots). Plants were well-watered for two weeks before water was completely withheld for 18 or 21 days. Three days after rewatering, survival rate was scored by counting plants that remained alive. During the entire period, all pots in the same flat were rotated frequently to minimize differences in the growth environment. Three independent experiments were performed each including nine pots or 81 individual plants.

### Leaf Infrared Thermography

To monitor the leaf temperature, thermal imaging was performed as described previously with slight modification (Zhao et al., 2016a). Well-watered four-week-old plants grown under ∼70% relative humidity were transferred to a low humidity condition (40% relative humidity). Three days thereafter, infrared energy emitted from the leaves was recorded using a VarioCAM HD infrared camera (InfraTec). Leaf temperature was deduced and temperature distribution images were generated using the FLIR Tools+ professional software. For each genotype, plants in four different pots were analyzed and similar results were obtained.

### Measurements of ABA Content

Endogenous ABA was extracted and measured using an ELISA Kit for Abscisic Acid (Cloud-Clone). Briefly, one gram of freshly harvested seedlings was homogenized in liquid nitrogen and extracted with 6 mL 80% methyl alcohol overnight at 4℃. The supernatant was collected and the pellet was further extracted with 2 mL 80% methyl alcohol and centrifuged at 2000*g*. The combined supernatant was reduced to a volume of 2 mL by evaporation on a rotary evaporator and then mixed with 1 mL petroleum ether to remove chlorophylls. The aqueous phase was collected and methyl alcohol was removed by evaporation. A quadruplicate of 50 μL aqueous sample was added into pre-coated 96-well plate provide in the kit, sequentially mixed with 50 μL detection reagent A, 100 μL detection reagent B, 90 μL substrate solution, and 50 μL stop solution as instructed by the manufacturer. Absorbance at 450 nm was read on a Cytation 5 multi-mode plate reader (BioTeK). Statistical analysis was performed on three experiments using individual batches of seedlings or plants with each experiment containing four technical repeats (Supplemental Data Set 1).

ABA was also purified and quantified in adult plants using the UPLC-MS/MS method as previously described with minor modifications (Fu et al. 2012). Briefly, one g of leaves from each genotype was collected and homogenized in liquid nitrogen. UPLC-MS/MS analysis was performed on an UltiMate 3000 UHPLC+ system (Thermo Fisher Scientific) coupled with a TSQ Quantiva triple quadrupole mass spectrometer (Thermo Fisher Scientific) at Wuhan Greensword Creation Technology. Statistical analysis was performed on three experiments using individual batches of leaves (Supplemental Data Set 3).

### Measurement of Anthocyanin Levels

Measurement of anthocyanin was performed as described previously (Zhang et al., 2014). Rosette leaves of phase III plants treated with or without drought stress for the indicated times were harvested, weighed, and homogenized in liquid nitrogen. One hundred mg homogenized sample was suspended in 0.3 mL of 1% HCl in methanol and incubated overnight at 4℃. Following the addition of 0.2 mL deionized water and extraction with an equal volume of chloroform, the aqueous phase was collected and subjected to spectrophotometric measurement. The quantity of anthocyanins was determined as A_530_ - 0.25(A_657_) and normalized to the fresh weight of each sample.

### Electrolyte Leakage Assay

Electrolyte leakage was measured as previously described (Zhao et al., 2016b). Briefly, 0.5 g rosette leaf was washed with double-distilled water and placed in 50 mL tubes filled with 20 mL double-distilled water. The electrical conductivity of this solution (L1) was measured with a LE703 conductivity meter (Mettler Toledo) after 1 h of gentle shaking (80 rpm) at room temperature. The samples were then boiled for 20 min and measured a second time for conductivity (L2). Electrolyte leakage was calculated as (L1/L2) × 100%.

### Promoter Activity Assays

Effects of SPL7 on target promoters were tested employing the REN/LUC dual luciferase system as described with minor modifications (Liu et al., 2014). The native *ZEP*, *NCED3*, and *AAO3* promoter regions were cloned from Arabidopsis genomic DNA into the modified p1305.1-LUC vector using primers listed in Supplemental Table 1 to generate the reporter constructs. The promoters of *MIR408* and *ACTIN* were also cloned into the p1305.1-LUC-35S-REN vector as controls. The coding region of *SPL7* and *SPL7^SBP^* were cloned into the pJIM19 vector to generate the *35S:SPL7-GFP* and *35S:SPL7^SBP^-GFP* effector constructs, respectively. The sequence-confirmed reporter and effector constructs were transformed into *Agrobacterium* strain GV3101, which was then used to co-infiltrate tobacco leaf epidermal cells. Two days after transfection, dual luciferase reactions were carried out using the Dual-Glo Luciferase Assay System (Promega) following the manufacturer’s protocol. Chemiluminescence was detected using a NightSHADE LB 985 system (Berthold) and quantified using a Multimode Reader LB 942 luminometer (Berthold). Replicates were individual transfection events.

### RT-qPCR

Total RNA was isolated using the Trizol reagent (Invitrogen), treated with DNase I (TaKaRa), and reverse transcribed using the PrimeScript II 1^st^ Strand cDNA Synthesis Kit (TaKaRa). qPCR was carried out using the SYBR Green master mix on a QuantStudio 3 Real-Time PCR System (Applied Biosystems) using primers listed in Supplemental Table 1. *ACTIN7* was used as the internal control and normalization standard. Quantification of differential expression was based on three replicate qPCR reactions performed on the same cDNA.

### ChIP-qPCR

ChIP experiments were performed as previously described (Zhou et al., 2018). Ten-day-old *35S:SPL7^SBP^-GFP* and wild type seedlings were fixed in phosphate buffer saline (pH 7.4) with 1% formaldehyde under vacuum for 10 min and then homogenized in liquid nitrogen. Extracted chromatin was sonicated to ∼200-500 bp at 4℃ using a Bioruptor (Diagenode). For immunoprecipitation, a GFP antibody (Abcam, catalog # ab290, 1:1000 (v/v) dilution) and a rabbit IgG antibody (Abcam, catalog # ab171870, 1:1000 (v/v) dilution) together with protein A-Sepharose beads (GE Healthcare) were used. After ChIP, equal amount of input and immunoprecipitated DNA was subjected to qPCR analysis using primers listed in Supplementary Table 1. Quantification was based on three replicate qPCR reactions performed on the same DNA.

### LUC Assay

The *SPL7_pro_:SPL7*, *SPL7_pro_:SPL7-LUC*, and *35S:LUC* seedlings were grown on half-strength MS medium supplemented with the indicated concentrations of Cu for ten days. The seedlings were then sprayed with 200 μg mL^-1^ potassium luciferin (Gold Biotech). Luciferase activity was quantified using a PyLoN2048B luminescence imaging system (Lumazone) with a 10 min exposure time. Three independent experiments were performed each including ten seedlings.

### Differential Gene Expression Analyses

RNA-sequencing data were obtained from NCBI BioProject (https://www.ncbi.nlm.nih.gov/bioproject/). The processed reads were directly used for transcript quantification using R scripts. Differentially expressed genes were called using fold change cutoff of ± 1.5 against the repective controls. The false discovery rate (FDR) was used for correcting multiple comparisons and a cutoff of 0.05 was applied. Clustering analysis and heatmap of gene expression profile were performed and visualized using R scripts. GO analysis was performed using AgriGO (http://bioinfo.cau.edu.cn/agriGO/) and visualized using R scripts.

### Statistical Analyses

Student’s *t*-test was used for comparing means between two samples. For all multiple comparisons, one-way ANOVA was first conducted to test the significance of the difference among different group means. Dunnett’s test was used post hoc for multiple comparisons between two group means at the indicated significance levels. Detailed statistical reports are shown in Supplemental Dataset 1.

### Phylogenetic Analysis

The ZEP, NCED, AAO related protein sequences for *A. thaliana*, *S. lycopersicum*, *O. sativa*, *Z. mays*, *A. trichopoda*, *Z. marina*, *S. moellendorffii*, *P. patens*, *M. polymorpha*, and *C. reinhardtii* were obtained from Phytozome v12 (https://phytozome.jgi.doe.gov) using the built-in search tool. The ZEP, NCED, AAO homologs in *G. biloba* and *K. flaccidum* were identified by all-against-all similarity searches using BLASTP with E-value cutoff set at e^-5^ from the respective genome annotations at ftp://parrot.genomics.cn/gigadb/pub/10.5524/100001_101000/100613/Ginkgo_biloba.HiC.protei n.fasta and http://www.plantmorphogenesis.bio.titech.ac.jp/cgi-bin/blast/blast_www_klebsormidium.cgi. Multiple sequence alignments were performed using the MUSCLE program in the MEGA X software with the default settings(Kumar et al., 2018). Alignments of the ZEP, NCED, and AAO related proteins are provided as Supplemental Files 1 to 3, respectively. Phylogenetic trees were constructed using MEGA X by applying the neighbor-joining method, 1000 bootstrap replicates, and the Jones-Taylor-Thornton model.

### Accession Number

Sequence data from this article can be found in the *Arabidopsis* Genome Initiative or GenBank/EMBL databases under the following accession numbers: *COR15B* (At2g42530), *HAB1* (At1g72770), *DREB2A* (AT5G05410), *SPL7* (At5g18830), *ZEP* (At5g67030), *NCED3* (At3g14440), *NCED6* (At3g24220), *NCED9* (At1g78390), *AAO3* (At2g27150), *ABI1* (At4g26080), *ABI5* (At2g36270), *ABF1* (At1g49720), *RD20* (At2g33380), *RD29A* (At5g52310), and *RD29B* (At5g52300). T-DNA insertion mutant used is *spl7-1* (SALK_093849C). RNA sequencing data can be found at the National Center for Biotechnology Information Sequence Read Archive under the accession numbers PRJNA350795, SRS1792515, and SRS1792516.

## Supplemental Data

**Supplemental Figure 1.** Generation and Characterization of *SPL7* Related Lines.

**Supplemental Figure 2.** SPL7 Regulates Plant Growth at Checkpoint II.

**Supplemental Figure 3.** The *spl7* Mutants Are Insensitive to High Cu Induced Growth Arrest.

**Supplemental Figure 4.** Expression Profiles of Representative Drought Inducible Genes.

**Supplemental Figure 5.** Characterization of LUC Reporter Lines.

**Supplemental Figure 6.** Analysis of the Luminescence Levels in Various Reporter Plants.

**Supplemental Figure 7.** Expression Profile of Excess Cu and SPL7 Co-regulated Genes.

**Supplemental Figure 8.** Cu and SPL7 Influence *NCED3* and *AAO3* Transcript Abundance.

**Supplemental Figure 9.** Comparison of the occurrence of the GTAC tetranucleotide in the *ZEP/FAD/NAD(P)-BINDING OXIDOREDUCTASE* promoters.

**Supplemental Figure 10.** Comparison of the occurrence of the GTAC tetranucleotide in the *NCED/CAROTENOID CLEAVAGE DIOXYGENASE (CCD)* promoters.

**Supplemental Figure 11.** Comparison of the occurrence of the GTAC tetranucleotide in the *AAO/XANTHINE DEHYDROGENASE (XDH)* promoters.

**Supplemental Table 1.** Oligonucleotide Sequences of the Primers Used in This Study.

**Supplemental Data Set 1.** Results of Statistical Analyses.

**Supplemental File 1.** Sequence Alignment of ZEP Homologs.

**Supplemental File 2.** Sequence Alignment of NCED Homologs.

**Supplemental File 3.** Sequence Alignment of AAO Homologs.

## ACKNOWLEDGEMENTS

We thank Drs. Yan Guo and Yue Zhou for advice on the infrared thermography and ChIP experiments. This work was supported by grants from the National Key Research and Development Program of China (2018YFE0204700) and the National Natural Science Foundation of China (31621001).

## AUTHOR CONTRIBUTIONS

L.L. designed and supervised the research. Y.Y., J.D., Y.T. C.H. and Z.L. performed the research. Y.Y., and Z.K. analyzed the data. Y.Y. and L.L. wrote the paper.

## REFERENCES

Abdel-Ghany, S.E. (2009). Contribution of plastocyanin isoforms to photosynthesis and copper homeostasis in Arabidopsis thaliana grown at different copper regimes. Planta 229, 767–779.

Abdel-Ghany, S.E., and Pilon, M. (2008). MicroRNA-mediated systemic down-regulation of copper protein expression in response to low copper availability in Arabidopsis. J Biol Chem 283, 15932–15945.

An, Z., Jing, W., Liu, Y., and Zhang, W. (2008). Hydrogen peroxide generated by copper amine oxidase is involved in abscisic acid-induced stomatal closure in Vicia faba. J Exp Bot 59, 815–825.

Basu, S., Ramegowda, V., Kumar, A., and Pereira, A. (2016). Plant adaptation to drought stress. F1000Res 5.

Bernal, M., Casero, D., Singh, V., Wilson, G.T., Grande, A., Yang, H., Dodani, S.C., Pellegrini, M., Huijser, P., Connolly, E.L., Merchant, S.S., and Krämer, U. (2012). Transcriptome sequencing identifies SPL7-regulated copper acquisition genes FRO4/FRO5 and the copper dependence of iron homeostasis in Arabidopsis. Plant Cell 24, 738–761.

Bowman, J.L., Kohchi, T., Yamato, K.T., Jenkins, J., Shu, S., Ishizaki, K., Yamaoka, S., Nishihama, R., Nakamura, Y., Berger, F., Adam, C., Aki, S.S., Althoff, F., Araki, T., Arteaga-Vazquez, M.A., Balasubrmanian, S., Barry, K., Bauer, D., Boehm, C.R., Briginshaw, L., Caballero-Perez, J., Catarino, B., Chen, F., Chiyoda, S., Chovatia, M., Davies, K.M., Delmans, M., Demura, T., Dierschke, T., Dolan, L., Dorantes-Acosta, A.E., Eklund, D.M., Florent, S.N., Flores-Sandoval, E., Fujiyama, A., Fukuzawa, H., Galik, B., Grimanelli, D., Grimwood, J., Grossniklaus, U., Hamada, T., Haseloff, J., Hetherington, A.J., Higo, A., Hirakawa, Y., Hundley, H.N., Ikeda, Y., Inoue, K., Inoue, S.I., Ishida, S., Jia, Q., Kakita, M., Kanazawa, T., Kawai, Y., Kawashima, T., Kennedy, M., Kinose, K., Kinoshita, T., Kohara, Y., Koide, E., Komatsu, K., Kopischke, S., Kubo, M., Kyozuka, J., Lagercrantz, U., Lin, S.S., Lindquist, E., Lipzen, A.M., Lu, C.W., De Luna, E., Martienssen, R.A., Minamino, N., Mizutani, M., Mizutani, M., Mochizuki, N., Monte, I., Mosher, R., Nagasaki, H., Nakagami, H., Naramoto, S., Nishitani, K., Ohtani, M., Okamoto, T., Okumura, M., Phillips, J., Pollak, B., Reinders, A., Rövekamp, M., Sano, R., Sawa, S., Schmid, M.W., Shirakawa, M., Solano, R., Spunde, A., Suetsugu, N., Sugano, S., Sugiyama, A., Sun, R., Suzuki, Y., Takenaka, M., Takezawa, D., Tomogane, H., Tsuzuki, M., Ueda, T., Umeda, M., Ward, J.M., Watanabe, Y., Yazaki, K., Yokoyama, R., Yoshitake, Y., Yotsui, I., Zachgo, S., and Schmutz, J. (2017). Insights into land plant evolution garnered fromthe Marchantia polymorpha genome. Cell 171, 287–304.e215.

Boyer, J.S. (1982). Plant productivity and environment. Science 218, 443–448.

Burkhead, J.L., Reynolds, K.A., Abdel-Ghany, S.E., Cohu, C.M., and Pilon, M. (2009). Copper homeostasis. New Phytol 182, 799–816.

Chen, K., Li, G.J., Bressan, R.A., Song, C.P., Zhu, J.K., and Zhao, Y. (2020). Abscisic acid dynamics, signaling, and functions in plants. J Integr Plant Biol 62, 25–54.

Claeys, H., and Inzé, D. (2013). The agony of choice: how plants balance growth and survival under water-limiting conditions. Plant Physiol 162, 1768–1779.

Clough, S.J., and Bent, A.F. (1998). Floral dip: a simplified method for Agrobacterium-mediated transformation of Arabidopsis thaliana. Plant J 16, 735–743.

Eklund, D.M., Kanei, M., Flores-Sandoval, E., Ishizaki, K., Nishihama, R., Kohchi, T., Lagercrantz, U., Bhalerao, R.P., Sakata, Y., and Bowman, J.L. (2018). An evolutionarily conserved abscisic acid signaling pathway regulates dormancy in the liverwort Marchantia polymorpha. Curr Biol 28, 3691–3699.e3693.

Endo, A., Sawada, Y., Takahashi, H., Okamoto, M., Ikegami, K., Koiwai, H., Seo, M., Toyomasu, T., Mitsuhashi, W., Shinozaki, K., Nakazono, M., Kamiya, Y., Koshiba, T., and Nambara, E. (2008). Drought induction of Arabidopsis 9-cis-epoxycarotenoid dioxygenase occurs in vascular parenchyma cells. Plant Physiol 147, 1984–1993.

Finney, L.A., and O’Halloran, T.V. (2003). Transition metal speciation in the cell: insights fromthe chemistry of metal ion receptors. Science 300, 931–936.

Fu, J., Chu, J., Sun, X., Wang, J., and Yan, C. (2012). Simple, rapid, and simultaneous assay of multiple carboxyl containing phytohormones in wounded tomatoes by UPLC-MS/MS using single SPE purification and isotope dilution. Anal Sci 28, 1081–1087.

Garcia-Molina, A., Xing, S., and Huijser, P. (2014). Aconserved KIN17 curved DNA-binding domain protein assembles with SQUAMOSA PROMOTER-BINDING PROTEIN-LIKE7 to adapt Arabidopsis growth and development to limiting copper availability. Plant Physiol 164, 828–840.

Gupta, A., Rico-Medina, A., and Caño-Delgado, A.I. (2020). The physiology of plant responses to drought. Science 368, 266–269.

Hauser, F., Waadt, R., and Schroeder, J.I. (2011). Evolution of abscisic acid synthesis and signaling mechanisms. Curr Biol 21, R346–355.

Iuchi, S., Kobayashi, M., Taji, T., Naramoto, M., Seki, M., Kato, T., Tabata, S., Kakubari, Y., Yamaguchi-Shinozaki, K., and Shinozaki, K. (2001). Regulation of drought tolerance by gene manipulation of 9-cis-epoxycarotenoid dioxygenase, a key enzyme in abscisic acid biosynthesis in Arabidopsis. Plant J 27, 325–333.

Jiang, A., Guo, Z., Pan, J., Yang, Y., Zhuang, Y., Zuo, D., Hao, C., Gao, Z., Xin, P., Chu, J., Zhong, S., and Li, L. (2021). The PIF1-miR408-PLANTACYANIN repression cascade regulates light-dependent seed germination. Plant Cell. Online ahead of print.

Khandelwal, A., Cho, S.H., Marella, H., Sakata, Y., Perroud, P.F., Pan, A., and Quatrano, R.S. (2010). Role of ABA and ABI3 in desiccation tolerance. Science 327, 546.

Komatsu, K., Takezawa, D., and Sakata, Y. (2020). Decoding ABA and osmostress signalling in plants froman evolutionary point of view. Plant Cell Environ 43, 2894–2911.

Kropat, J., Tottey, S., Birkenbihl, R.P., Depège, N., Huijser, P., and Merchant, S. (2005). Aregulator of nutritional copper signaling in Chlamydomonas is an SBP domain protein that recognizes the GTAC core of copper response element. Proc Natl Acad Sci U S A 102, 18730–18735.

Kumar, S., Stecher, G., Li, M., Knyaz, C., and Tamura, K. (2018). MEGAX: molecular evolutionary genetics analysis across computing platforms. Mol Biol Evol 35, 1547–1549.

Kuper, J., Llamas, A., Hecht, H.J., Mendel, R.R., and Schwarz, G. (2004). Structure of the molybdopterin-bound Cnx1G domain links molybdenum and copper metabolism. Nature 430, 803–806.

Li, P., Li, Y.J., Zhang, F.J., Zhang, G.Z., Jiang, X.Y., Yu, H.M., and Hou, B.K. (2017). The Arabidopsis UDP-glycosyltransferases UGT79B2 and UGT79B3, contribute to cold, salt and drought stress tolerance via modulating anthocyanin accumulation. Plant J 89, 85–103.

Liu, Q., Wang, F., and Axtell, M.J. (2014). Analysis of complementarity requirements for plant microRNA targeting using a Nicotiana benthamiana quantitative transient assay. Plant Cell 26, 741–753.

Liu, Z., Yan, J.P., Li, D.K., Luo, Q., Yan, Q., Liu, Z.B., Ye, L.M., Wang, J.M., Li, X.F., and Yang, Y. (2015). UDP-glucosyltransferase71c5, a major glucosyltransferase, mediates abscisic acid homeostasis in Arabidopsis. Plant Physiol 167, 1659–1670.

McAdam, S.A.M., and Brodribb, T.J. (2018). Mesophyll Cells Are the Main Site of Abscisic Acid Biosynthesis in Water-Stressed Leaves. Plant Physiol 177, 911–917.

McAinsh, M.R., Brownlee, C., and Hetherington, A.M. (1990). Abscisic acid-induced elevation of guard cell cytosolic Ca2+ precedes stomatal closure. Nature 343, 186–188.

Molina-Heredia, F.P., Wastl, J., Navarro, J.A., Bendall, D.S., Hervás, M., Howe, C.J., and De La Rosa, M.A. (2003). Photosynthesis: a new function for an old cytochrome? Nature 424, 33–34.

Murashige, T., Skoog, F. (1962) Arevised medium for rapid growth and bio assays with tobacco tissue cultures. Physiol Plant 15, 473–497.

Nakabayashi, R., Yonekura-Sakakibara, K., Urano, K., Suzuki, M., Yamada, Y., Nishizawa, T., Matsuda, F., Kojima, M., Sakakibara, H., Shinozaki, K., Michael, A.J., Tohge, T., Yamazaki, M., and Saito, K. (2014). Enhancement of oxidative and drought tolerance in Arabidopsis by overaccumulation of antioxidant flavonoids. Plant J 77, 367–379.

Nambara, E., and Marion-Poll, A. (2005). Abscisic acid biosynthesis and catabolism. Annu Rev Plant Biol 56, 165–185.

Pan, J., Huang, D., Guo, Z., Kuang, Z., Zhang, H., Xie, X., Ma, Z., Gao, S., Lerdau, M.T., Chu, C., and Li, L. (2018). Overexpression of microRNA408 enhances photosynthesis, growth, and seed yield in diverse plants. J Integr Plant Biol 60, 323–340.

Pätsikkä, E., Kairavuo, M., Sersen, F., Aro, E.M., and Tyystjärvi, E. (2002). Excess copper predisposes photosystem II to photoinhibition in vivo by outcompeting iron and causing decrease in leaf chlorophyll. Plant Physiol 129, 1359–1367.

Pei, Z.M., Murata, Y., Benning, G., Thomine, S., Klüsener, B., Allen, G.J., Grill, E., and Schroeder, J.I. (2000). Calciumchannels activated by hydrogen peroxide mediate abscisic acid signalling in guard cells. Nature 406, 731–734.

Peñarrubia, L., Romero, P., Carrió-Seguí, A., Andrés-Bordería, A., Moreno, J., and Sanz, A. (2015). Temporal aspects of copper homeostasis and its crosstalk with hormones. Front Plant Sci 6, 255.

Pope, C.R., Flores, A.G., Kaplan, J.H., and Unger, V.M. (2012). Structure and function of copper uptake transporters. Curr Top Membr 69, 97–112.

Pourcel, L., Routaboul, J.M., Cheynier, V., Lepiniec, L., and Debeaujon, I. (2007). Flavonoid oxidation in plants: frombiochemical properties to physiological functions. Trends Plant Sci 12, 29–36.

Puig, S., and Thiele, D.J. (2002). Molecular mechanisms of copper uptake and distribution. Current Opinion in Chemical Biology 6, 171–180.

Rae, T.D., Schmidt, P.J., Pufahl, R.A., Culotta, V.C., and O’Halloran, T.V. (1999). Undetectable intracellular free copper: the requirement of a copper chaperone for superoxide dismutase. Science 284, 805–808.

Reyt, G., Chao, Z., Flis, P., Salas-González, I., Castrillo, G., Chao, D.Y., and Salt, D.E. (2020). Uclacyanin proteins are required for lignified nanodomain formation within casparian strips. Curr Biol 30, 4103–4111.e4106.

Robinson, N.J., and Winge, D.R. (2010). Copper metallochaperones. Annu. Rev. Biochem. 79, 537–562.

Rodríguez, F.I., Esch, J.J., Hall, A.E., Binder, B.M., Schaller, G.E., and Bleecker, A.B. (1999). Acopper cofactor for the ethylene receptor ETR1 from Arabidopsis. Science 283, 996–998.

Ross, G.S., Elder, P.A., McWha, J.A., Pearce, D., and Pharis, R.P. (1987). The development of an indirect enzyme linked immunoassay for abscisic Acid. Plant Physiol 85, 46–50.

Sekimoto, H., Seo, M., Kawakami, N., Komano, T., Desloire, S., Liotenberg, S., Marion-Poll, A., Caboche, M., Kamiya, Y., and Koshiba, T. (1998). Molecular cloning and characterization of aldehyde oxidases in Arabidopsis thaliana. Plant Cell Physiol 39, 433–442.

Seo, M., Peeters, A.J., Koiwai, H., Oritani, T., Marion-Poll, A., Zeevaart, J.A., Koornneef, M., Kamiya, Y., and Koshiba, T. (2000). The Arabidopsis aldehyde oxidase 3 (AAO3) gene product catalyzes the final step in abscisic acid biosynthesis in leaves. Proc Natl Acad Sci U S A 97, 12908–12913.

Sun, Y., Harpazi, B., Wijerathna-Yapa, A., Merilo, E., de Vries, J., Michaeli, D., Gal, M., Cuming, A.C., Kollist, H., and Mosquna, A. (2019). A ligand-independent origin of abscisic acid perception. Proc Natl Acad Sci U S A 116, 24892–24899.

Takezawa, D., Komatsu, K., and Sakata, Y. (2011). ABA in bryophytes: how a universal growth regulator in life became a plant hormone? J Plant Res 124, 437–453.

Takezawa, D., Watanabe, N., Ghosh, T.K., Saruhashi, M., Suzuki, A., Ishiyama, K., Somemiya, S., Kobayashi, M., and Sakata, Y. (2015). Epoxycarotenoid-mediated synthesis of abscisic acid in Physcomitrella patens implicating conserved mechanisms for acclimation to hyperosmosis in embryophytes. New Phytol 206, 209–219.

Umezawa, T., Nakashima, K., Miyakawa, T., Kuromori, T., Tanokura, M., Shinozaki, K., and Yamaguchi-Shinozaki, K. (2010). Molecular basis of the core regulatory network in ABA responses: sensing, signaling and transport. Plant Cell Physiol 51, 1821–1839.

Vishwakarma, K., Upadhyay, N., Kumar, N., Yadav, G., Singh, J., Mishra, R.K., Kumar, V., Verma, R., Upadhyay, R.G., Pandey, M., and Sharma, S. (2017). Abscisic acid signaling and abiotic stress tolerance in plants: a review on current knowledge and future prospects. Front Plant Sci 8, 161.

Wang, K., He, J., Zhao, Y., Wu, T., Zhou, X., Ding, Y., Kong, L., Wang, X., Wang, Y., Li, J., Song, C.P., Wang, B., Yang, S., Zhu, J.K., and Gong, Z. (2018a). EAR1 negatively regulates ABA signaling by enhancing 2C protein phosphatase activity. Plant Cell 30, 815–834.

Wang, S., Saito, T., Ohkawa, K., Ohara, H., Suktawee, S., Ikeura, H., and Kondo, S. (2018b). Abscisic acid is involved in aromatic ester biosynthesis related with ethylene in green apples. J Plant Physiol 221, 85–93.

Weigel, M., Varotto, C., Pesaresi, P., Finazzi, G., Rappaport, F., Salamini, F., and Leister, D. (2003). Plastocyanin is indispensable for photosynthetic electron flow in Arabidopsis thaliana. J Biol Chem 278, 31286–31289.

Xiong, L., and Zhu, J.K. (2003). Regulation of abscisic acid biosynthesis. Plant Physiol 133, 29–36.

Xiong, L., Ishitani, M., Lee, H., and Zhu, J.K. (2001). The Arabidopsis LOS5/ABA3 locus encodes a molybdenum cofactor sulfurase and modulates cold stress- and osmotic stress-responsive gene expression. Plant Cell 13, 2063–2083.

Xiong, L., Lee, H., Ishitani, M., and Zhu, J.K. (2002). Regulation of osmotic stress-responsive gene expression by the LOS6/ABA1 locus in Arabidopsis. J Biol Chem 277, 8588–8596.

Xu, Z.Y., Lee, K.H., Dong, T., Jeong, J.C., Jin, J.B., Kanno, Y., Kim, D.H., Kim, S.Y., Seo, M., Bressan, R.A., Yun, D.J., and Hwang, I. (2012). Avacuolar β-glucosidase homolog that possesses glucose-conjugated abscisic acid hydrolyzing activity plays an important role in osmotic stress responses in Arabidopsis. Plant Cell 24, 2184–2199.

Yamasaki, H., Hayashi, M., Fukazawa, M., Kobayashi, Y., and Shikanai, T. (2009). SQUAMOSA Promoter Binding Protein-Like7 is a central regulator for copper homeostasis in Arabidopsis. Plant Cell 21, 347–361.

Yan, J., Chia, J.C., Sheng, H., Jung, H.I., Zavodna, T.O., Zhang, L., Huang, R., Jiao, C., Craft, E.J., Fei, Z., Kochian, L.V., and Vatamaniuk, O.K. (2017). Arabidopsis pollen fertility requires the transcription factors CITF1 and SPL7 that regulate copper delivery to anthers and jasmonic acid synthesis. Plant Cell 29, 3012–3029.

Ye, N., Li, H., Zhu, G., Liu, Y., Liu, R., Xu, W., Jing, Y., Peng, X., and Zhang, J. (2014). Copper suppresses abscisic acid catabolismand catalase activity, and inhibits seed germination of rice. Plant Cell Physiol 55, 2008–2016.

Zhang, H., and Li, L. (2013). SQUAMOSA promoter binding protein-like7 regulated microRNA408 is required for vegetative development in Arabidopsis. Plant J 74, 98–109.

Zhang, H., Zhao, Y., and Zhu, J.K. (2020). Thriving under stress: how plants balance growth and the stress response. Dev Cell 55, 529–543.

Zhang, H., Zhao, X., Li, J., Cai, H., Deng, X.W., and Li, L. (2014). MicroRNA408 is critical for the HY5-SPL7 gene network that mediates the coordinated response to light and copper. Plant Cell 26, 4933–4953.

Zhao, Q., Nakashima, J., Chen, F., Yin, Y., Fu, C., Yun, J., Shao, H., Wang, X., Wang, Z.Y., and Dixon, R.A. (2013). Laccase is necessary and nonredundant with peroxidase for lignin polymerization during vascular development in Arabidopsis. Plant Cell 25, 3976–3987.

Zhao, S., Jiang, Y., Zhao, Y., Huang, S., Yuan, M., Zhao, Y., and Guo, Y. (2016a). CASEIN KINASE1-LIKE PROTEIN2 regulates actin filament stability and stomatal closure via phosphorylation of actin depolymerizing factor. Plant Cell 28, 1422–1439.

Zhao, Y., Lin, S., Qiu, Z., Cao, D., Wen, J., Deng, X., Wang, X., Lin, J., and Li, X. (2015). MicroRNA857 is involved in the regulation of secondary growth of vascular tissues in Arabidopsis. Plant Physiol 169, 2539–2552.

Zhao, Y., Chan, Z., Gao, J., Xing, L., Cao, M., Yu, C., Hu, Y., You, J., Shi, H., Zhu, Y., Gong, Y., Mu, Z., Wang, H., Deng, X., Wang, P., Bressan, R.A., and Zhu, J.K. (2016b). ABA receptor PYL9 promotes drought resistance and leaf senescence. Proc Natl Acad Sci U S A 113, 1949–1954.

Zhou, Y., Wang, Y., Krause, K., Yang, T., Dongus, J.A., Zhang, Y., and Turck, F. (2018). Telobox motifs recruit CLF/SWN-PRC2 for H3K27me3 deposition via TRB factors in Arabidopsis. Nat Genet 50, 638–644.

Zhu, Y., Huang, P., Guo, P., Chong, L., Yu, G., Sun, X., Hu, T., Li, Y., Hsu, C.C., Tang, K., Zhou, Y., Zhao, C., Gao, W., Tao, W.A., Mengiste, T., and Zhu, J.K. (2020). CDK8 is associated with RAP2.6 and SnRK2.6 and positively modulates abscisic acid signaling and drought response in Arabidopsis. New Phytol 228, 1573–1590.

Zhuang, Y., and Li, L. (2020). Are cuproproteins part of the multi-protein framework for making the Casparian strip? Plant Signal Behav 15, 1798605.

Zhuang, Y., Zuo, D., Tao, Y., Cai, H., and Li, L. (2020). Laccase3-based extracellular domain provides possible positional information for directing Casparian strip formation in Arabidopsis. Proc Natl Acad Sci U S A 117, 15400–15402.

